# Global dynamics of microbial communities emerge from local interaction rules

**DOI:** 10.1101/2020.07.25.220822

**Authors:** Simon van Vliet, Christoph Hauert, Martin Ackermann, Alma Dal Co

## Abstract

Interactions between cells drive biological processes across all of life, from microbes in the environment to cells in multicellular organisms. Interactions often arise in spatially structured settings, where cells mostly interact with their neighbors. A central question is how local interactions shape the properties of biological systems. This question is very relevant in the context of microbial communities, such as biofilms, where cells live close by in space and are connected via a dense network of biochemical interactions. To understand and control the functioning of these communities, it is essential to uncover how community-level properties, such as the community composition, spatial arrangement, and growth rate, arise from these interactions. Here, we develop a mathematical framework that can predict community-level properties from the molecular mechanisms underlying the cell-cell interactions for systems consisting of two cell types. Our predictions can qualitatively reproduce measurements from an experimental cross-feeding community. For these cross-feeding communities, the community growth rate is reduced when cells interact only with few neighbors; as a result, some communities can co-exist in a well-mixed system, where cells can interact with all other cells, but not in systems where cells can interact only with close by neighbors. In general, our framework shows that key molecular parameters underlying the cell-cell interactions (e.g. the uptake and leakage rates of molecules) determine community-level properties. Our framework can be extended to a variety of systems of two interacting cell types, within and beyond the microbial world, and contributes to our understanding of how community-level dynamics and biological functions emerge from microscopic interactions between single cells.

## Introduction

Biological interactions pervade all of life. Interactions at lower levels of organization can confer new functionality at higher levels. For example, interactions between different cell types determine the functioning of organs and tissues in multicellular organisms and interactions between different species determine the processes an ecosystem performs [1, 2]. In natural systems, interactions often arise in spatially structured settings, where individual entities interact mostly with others that are close by in space [3]. When interactions are local, the spatial organization of the different entities defines their network of interaction. A central question is how the properties of biological systems emerge from this network of interactions. This has primarily been studied in the context of multicellular organisms, however it is also particularly relevant in the context of microbial communities [4, 5].

Microbial communities perform essential processes on our planet, and these processes often arise from interactions between species [6, 7]. Most microbial communities form spatially structured biofilms, where cells are embedded in an extracellular polymeric matrix that limits their movement [8]. In these communities, cells modify their local environment by secreting and taking up chemical compounds, and thus cells influence their neighbors’ growth, survival, and metabolic activity [9–16]. Most interactions are mediated by diffusible molecules and their strength decays with the distance between two interacting cells [17–20]. As a result, cells only interact within a limited distance, and the spatial organization of cells within the community determines which interactions are realized.

To predict and control the functioning of microbial systems, we need to uncover how cells interact in space and how these interactions determine community-level properties, such as the community composition, spatial arrangement, and growth rate [21]. Recent studies have made progress in this direction by characterizing the spatial arrangement of cells [10, 22, 23], the range over which cells interact [10, 11, 24, 25], how the sign of the interaction affects the spatial arrangement of the community [26,27], and how the spatial arrangement of cells affects community properties, such as their growth and species composition [5, 19, 22, 28–30].

Despite these recent advances, we do not understand well how local interaction rules scale up to determine community-level properties. In previous work, we demonstrated that local interaction rules can be measured in synthetic microbial communities and that mechanistic modeling can elucidate how these local interaction rules arise from molecular exchanges between cells [11]. Specifically, we focused on cells that exchange amino acids, which occurs commonly in nature [31]. We assembled communities of two cell types, where each type could not produce an amino acid and could only grow in mixed communities by receiving this amino acid from the other type. We showed that the two cell types interact within a small interaction range; moreover, we showed that the growth rate of a cell increased with the fraction of the partner type within the cell’s interaction range. The interaction range and the maximum growth rate can be different for the two cell types, and describe the local interaction rules between cells. These local interaction rules are modulated by biophysical parameters of the underlying molecular exchange: the interaction range is mostly set by the uptake rate of the exchanged molecules, while the maximum growth rate is set by the leakage rate of these molecules. Thus, the local interaction rules between cells arise from molecular-level parameters.

In the current work we demonstrate that the local interaction rules can predict community-level properties. We develop a mathematical framework which takes as input the local interaction rules and predicts community-level properties, such as the equilibrium frequency of cell types, their degree of spatial clustering, and the community growth rate. We experimentally validate our model’s predictions with the microbial community described above, and show that we can quantitatively predict the equilibrium ratio of the community, and can qualitatively predict the effect of the interaction range on the degree of spatial clustering. Our results suggests that, at least for simple biological systems, it is possible to scale up from molecular-level properties, to individual-level properties, to community-level properties. Properties at each level can be predicted from a few key quantities of the level below (Fig 1). These findings further our understanding of how scales are connected in biological systems.

**Fig 1:**
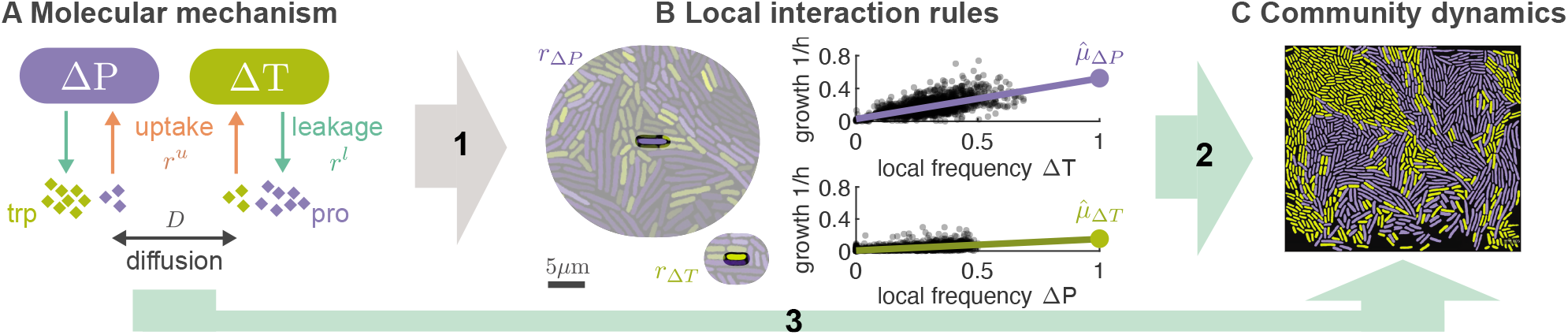
A mathematical framework to scale up from molecular-level properties, to individual-level properties, to community-level properties. We previously measured the local interaction rules for a cross-feeding community (B) and showed that these can be derived (arrow 1) from the molecular mechanisms of the interaction (A). Here we developed a mathematical model that can predict community-level properties (C) either from measured local rules (arrow 2) or directly from the underlying molecular mechanisms (arrow 3). (A) The community consists of two types of *Escherichia coli*: Δ*P* is unable to produce the amino acid proline and Δ*T* is unable to produce the amino acid tryptophan. Cells exchange amino acids with the environment through active uptake (with rate *r^u^*) and passive leakage (with rate *r^l^*). Amino acids are exchanged between cells through diffusion in the environment (with rate *D*). All rates differ between the two amino acids. (B) Local interaction rules can be fully described by two fundamental quantities: the size of the interaction neighborhood (*r*_Δ*T*_, *r*_Δ*P*_, left); and the growth function of a cell (characterized by 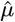, right). Each dot corresponds to the measured growth rate of a single cell, *n* = 2610 for ΔP and *n* = 2162 for ΔT, the line shows the result of a linear regression, data reproduced from [11]. (C) Our framework makes predictions for the community-level properties, such as the equilibrium frequency of the two types, their spatial arrangement, and growth rate.

## Model

### Local interaction rules

We present a mathematical framework that predicts community-level properties from local interaction rules for a variety of systems consisting of two interacting cell types. As a case study we focus on cross-feeding interactions, however our model applies more generally to any interactions that affect the growth rate of cells, and we discuss some of these possibilities later on. In previous work, we measured local interactions among cells in cross-feeding communities growing in two-dimensional environments [11]. We showed that two key quantities describe how cells interact in these communities: the interaction range (i.e. the size of the neighborhood a cell interacts with), and the maximum growth rate that cells achieves when all its neighbors are of the partner type (Fig 1B). We showed that the two cell types had different interaction ranges and maximum growth rates, and that these quantities were modulated by molecular parameters that were specific to the exchanged compounds.

Here, we build on this previous work and extend the biophysical model of cross-feeding communities to three dimensions, which is the more natural and general case (Fig S1, see Supporting information). In this model, cells live in dense three dimensional communities and exchange two compounds through leakage and uptake from the extracellular environment. We show that the local interaction rules between cells can be derived mathematically from key parameters (arrow 1 in Fig 1 [11], see Supporting information). The interaction range and the maximum growth rate are tuned independently: the interaction range primarily depends on the cell density and on the ratio between the uptake and diffusion rate of the molecules (Methods, Eq. 4); the maximum growth rate primarily depends on the leakage rate of the molecules (Eq. 6). Uptake, leakage, and diffusion rates are molecule specific; as a result the interaction range and maximum growth rate generally vary depending on which molecules are exchanged, as we also observed for our experimental community (Fig 1B).

A biophysical model is useful to understand how local interaction rules, like the interaction range, vary with molecular parameters, such as the uptake rate of exchanged compounds. However, the biophysical model is too complex to derive analytical expressions for the community-level dynamics, like the growth rate of the cross-feeding community or the fraction of cell types. Such analytical expressions are useful to understand how the molecular mechanisms of interaction between cells affect community dynamics. Reaching this goal is crucial if we want to control and engineer interactions in microbial systems.

To reach this goal, we here present a different mathematical framework that is suited to directly, and analytically, predict the community-level dynamics. We use the local interaction rules as input of the model and predict community-level properties, such as as the equilibrium frequency of the two types, their spatial arrangement, and the community growth rate (arrow 2 in Fig 1). The model connects the dynamics of molecules at a smaller spatial scale to the dynamics of the communities at a larger spatial scale (arrow 3 in Fig 1). The connection relies on our ability to reduce multiple biochemical parameters to a few key combinations, namely the interaction range and the maximum growth rate, that are the real drivers of community dynamics.

### Predicting community dynamics

Our framework generalizes previous models of interacting agents based on evolutionary graph theory [32–34]. We consider a spatial system of two cell types, A and B, that exchange essential cellular building blocks (Fig 2A). We assume that the total number of cells is constant and that each cell interacts with a fixed number of neighbors. In contrast to previous models, we allow the two cell types to interact with a different number of neighbors, as suggested by our experimental observations [11]. As a result, several simplifications no longer hold and extensions to previous work is required, which we discuss in the Supporting information. We assume that type A interacts with *r_A_* neighbors and that type B interacts with *r_B_* neighbors. We call *r_A_* and *r_B_* the *neighborhood sizes* and set *r_B_* ≥ *r_A_* (Fig 2B). Which cells interact with each other is encoded in the structure of the graph: all cells that can exchange building blocks are connected by links. To allow for different neighborhood sizes, we need to use directed graphs, instead of the undirected graphs. We indicate that a cell B is linked to a cell A with B←A; this link means that the focal cell B receives building blocks from the neighbor cell A. The neighbor cell A might not have a link to the focal cell B, because cells of type A receive building blocks from a smaller distance than cells of type B. Thus links are one-way, i.e., directed.

**Fig 2:**
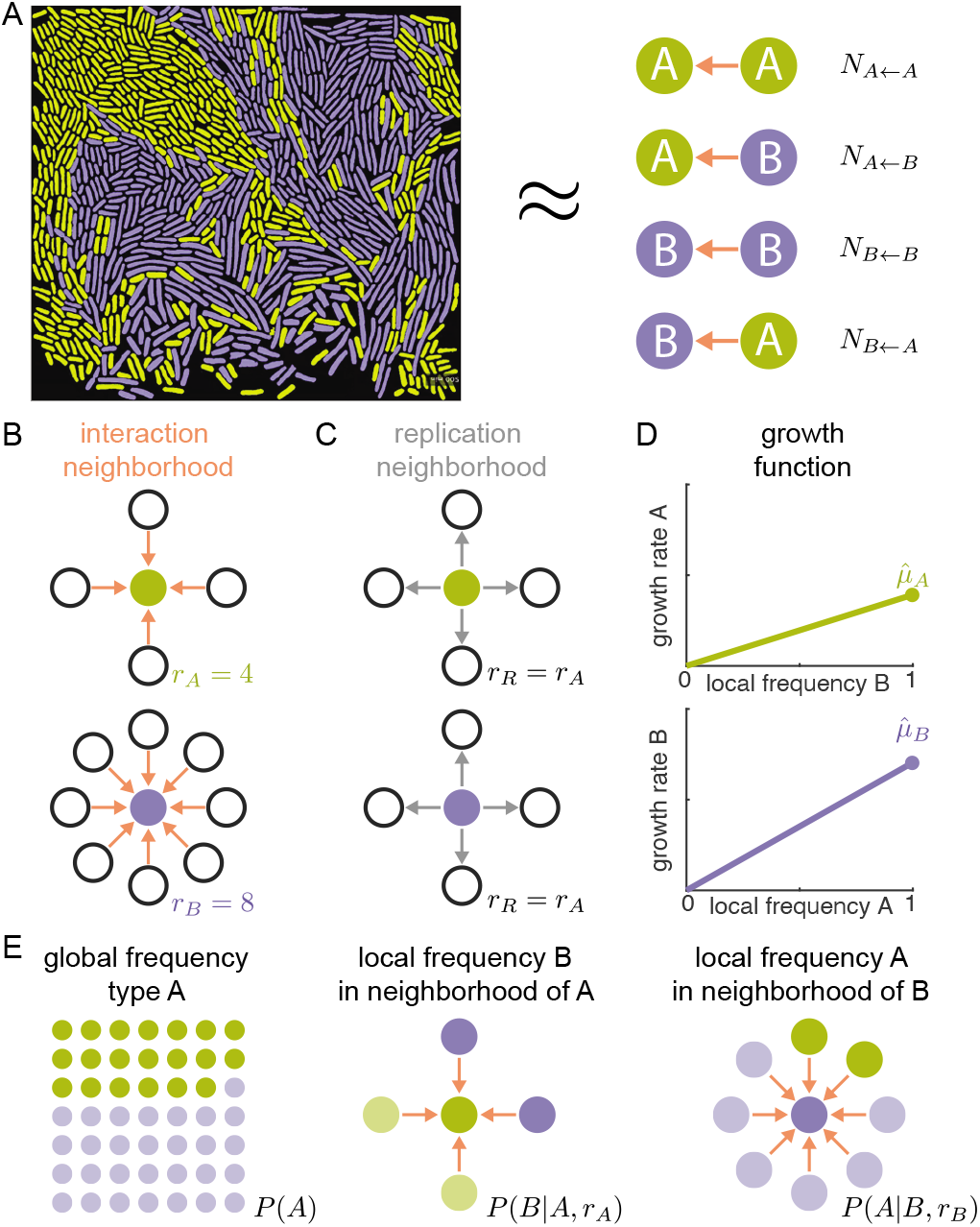
Pair approximation allows for a quantitative description of spatial systems. (A) Pair approximation assumes that a system of entities living in space and interacting with others close by (left side) can be fully described by tracking the number of all pairwise links between entities (right side). For example, a link counted in *N_X←Y_* indicates that focal cell X interacts with its neighbor Y. (B-D) A system is fully described by: (B) the interaction neighborhoods of both types, characterized by the neighborhood sizes *r_A_* and *r_B_*; (C) the replication neighborhood, assumed to be identical to the smallest interaction neighborhood; and (D) the growth functions of both types, characterized by the maximum growth rates 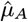 and 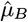. (E) Pair approximation has three dynamical variables that describe the global composition, *P* (*A*), and local composition *P* (*B*|*A, r_A_*) and *P* (*A*|*B, r_B_*) of the system.

Each cell (of type A or B) interacts with all (*r_A_* or *r_B_*) cells in its interaction neighborhood and these interactions determine its growth rate. We assume that the growth rate of a cell increases linearly with the frequency of the partner type within its interaction neighborhood (Fig 2D), as observed in the experimental communities [11]. Each type achieves its *maximum growth rate*, 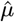, when it is completely surrounded by the partner type, and the maximum growth rate is different for the two types (Fig 2D). When a cell divides, it replaces a random neighbor within the replication neighborhood. We assume that the replication neighborhood is identical to the smaller interaction neighborhood (*r_R_* = *r_A_*, Fig 2C). This assumption has its limitations: in reality neighboring cells are not replaced, but rather pushed aside. Moreover, the replication neighborhood could be different from the smaller interaction neighborhood. However, this assumption does capture one of the most essential features of real biological systems: that cells place their offspring close by in space and thus become surrounded by their own type (see Supporting information for a more detailed discussion). This assumption does not strongly affect our quantitative predictions (see Fig S2), however it makes the model analytically tractable.

We implemented our model in two complementary ways: we used a cellular automaton to simulate the dynamics numerically (see Methods) and we used pair approximation to derive analytical predictions. Pair approximation assumes that the dynamics of a spatial system can be described by only specifying how often the different cell types are found in each others neighborhoods (Fig 2A) [32, 35, 36]. In our case, this means that the outcome of an interaction between a focal individual and one of its neighbors depends only on the identity of these two cells and not on the wider context in which these two cells are found. This approximation allows us to describe the dynamics of a spatial community (Fig 2A left) by only tracking how often cells of the different types are found within each others interaction neighborhood, i.e. by tracking the number of A←A, A←B, B←A, and B←B links (X←Y indicates that focal cell X receives building blocks from neighbor Y, Fig 2A right). Our model therefore generalizes previous models, where interactions were assumed to be symmetric and links undirected [32].

With these assumptions, we obtain the dynamical equations for the number of pairwise links in time (see Supporting information). Because the total number of cells is constant, we only have three independent variables (e.g. the number of A←A, A←B, B←A links) and we can express the fourth one (the number of B←B links) as a function of the other three.

Instead of tracking the number of links directly, it is convenient to change variables to *P* (*A*), the global probability of finding a type A cell; *P* (*B*|*A, r_A_*), the conditional probability of finding a type B cell in the interaction neighborhood of a type A focal cell; and *P* (*A*|*B, r_B_*), the conditional probability of finding a type A cell in the interaction neighborhood of a type B focal cell (Fig 2E). The first variable, *P* (*A*), characterizes the global composition (i.e. the frequency of type A) of the community. The other two variables characterize the local composition of the community by specifying the average frequency of the partner type within the interaction neighborhood of a focal cell. All other probabilities can be calculated from these three variables, e.g. the global frequency of type B *P* (*B*) is given by 1 − *P* (*A*). Moreover, the average growth rate of a cell in a spatial system can be calculated using the average local composition, i.e. the average frequency of the partner type within the cell’s interaction neighborhood.

### Steady state community properties

Pair approximation allows us to predict global properties of the community from the local interaction rules between cells. From the neighborhood sizes (*r_A_* and *r_B_*) and the maximum growth rates (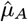 and 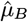, Fig 2B & D), we obtain a system of dynamical equations describing the global, *P* (*A*), and local, *P* (*B*|*A, r_A_*) and *P* (*A*|*B, r_B_*), composition of the system (Fig 2E). By solving the dynamical equation for steady state, we find analytical expressions for the global and local compositions at equilibrium (see Supporting information).

Pair approximation predicts that the global composition of the community reaches a steady state in which the frequency of type A is given by:

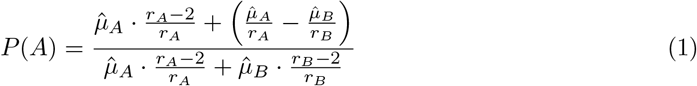

When the neighborhood size is large (*r_A_, r_B_* ≫ 1), this equation simplifies to the predicted frequency in a well-mixed system (see Supporting information):

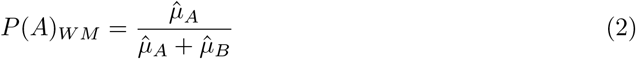

The same global community composition can correspond to many different local spatial arrangements of the two cell types. For example, the two cell types could be highly mixed in space or completely segregated. The spatial arrangement matters because cells interact only locally: the growth of a cell depends on the local frequency of the partner type, *P* (*B*|*A, r_A_*) and *P* (*A*|*B, r_B_*), which can be different from the global frequency, *P* (*B*) and *P* (*A*), when the cell types are clustered in space. A relevant quantity to describe a spatial system is thus the ratio between the local and global frequency of the partner type, and pair approximation predicts that at steady state:

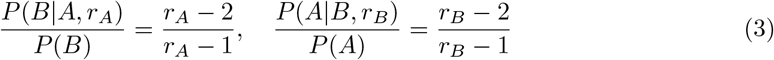

This equation shows that the frequency of the partner type in the interaction neighborhood is lower than one would expect from the global composition of the community. This happens because cells place offspring close to themselves when they divide. As a consequence, the local frequency of the partner type as much as 50% when the neighborhood size is small (e.g., when cells interact with three neighbors). One the other hand, when the neighborhood size is large, the local frequency of the partner type approaches the global frequency. Moreover, the dimensionality of the system matters: given the same interaction range (*R*), cells growing in two dimensional sheets have fewer neighbors (α *R*^2^) compared to cells growing in three dimensional structures (α *R*^3^). The interaction range (i.e. the distance over which molecules are exchanged) is similar in two and three dimensions because it mostly depends on the ratio of uptake and diffusion rate (see Supporting information). However the number of cells within the interaction range varies in two and three dimensions: cells in two-dimensional systems have fewer neighbors than cells in three-dimensional systems, which results in a stronger reduction of the local frequency of the partner type, and thus in slower growth.

## Results

### Local cell-cell interactions predict community properties

Our mathematical framework can predict the dynamics of cross-feeding communities when the local interaction rules between the two cell types are known. The model predicts that the equilibrium composition of the community is mostly set by the maximum growth rates of the two types. In general, the type with highest maximum growth rate constitutes the majority (Fig 3A). Also the neighborhood size affects the composition of the community: if the interaction neighborhoods are small, the composition shifts to the type with the faster maximum growth rate (Fig 3A)). Increasing the neighborhood size of even a single type moves the composition closer to the expected composition of a well-mixed system (Fig 3B, colored lines). The neighborhood sizes do not affect the community composition when both cell types have the same maximum growth rate (Fig 3B, black line). However, the neighborhood size affects the average composition of a cells’ interaction neighborhood in all cases: as the neighborhood size becomes smaller, the partner frequency in the interaction neighborhood decreases compared to the global frequency (Fig 3C). As a result, the average growth rate of the community decreases relative to an equivalent well-mixed community (Fig 3D).

**Fig 3:**
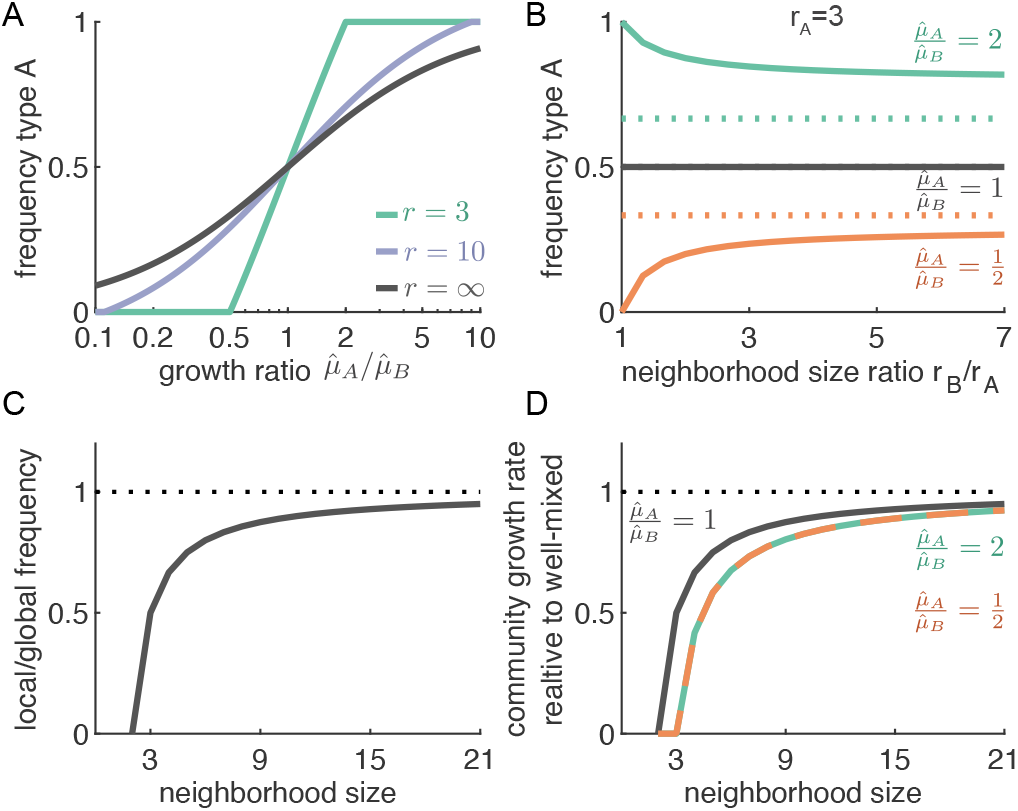
Community-level properties depend on local interaction rules. (A) The global composition of the community (global frequency of A, given by Eq. 1) primarily depends on the ratio of maximum growth rates of the two types. If the neighborhood sizes of both types are large (*r_A_* = *r_B_* = 10, purple line), the equilibrium frequency approaches that of a well-mixed system (black line, given by Eq. 2). If the neighborhood sizes are small (*r_A_* = *r_B_* = 3, green line), the type with the higher growth rate attains a higher frequency than in a well-mixed system. (B) The neighborhood size has only a minor effect on the global composition of the the community. The solid lines show the frequency of type A for spatial systems where *r_A_* is held constant at 3, while *r_B_* is increased. The equilibrium frequency varies as *r_B_* increases: for larger *r_B_*, the frequency in a spatial system (solid lines) moves closer to the frequency in a well-mixed system (dotted lines). This result holds when the types have different maximum growth rates (red and green curves) but not when they have equal maximum growth rates (black line). (C) The neighborhood size strongly affects the local composition of the community. Here the two types have the same maximum growth rate and neighborhood size. For both types, the local frequency of the partner cells is much lower than the global frequency, when the neighborhood size is small. (D) The neighborhood size strongly affects community growth rate. The average growth rate of cells is smaller when they have smaller interaction neighborhoods, because the local frequency of the partner around each cell is lower. This effect is stronger in communities where the types have different maximum growth (the red and green curves are below the black curve).

**Fig 4:**
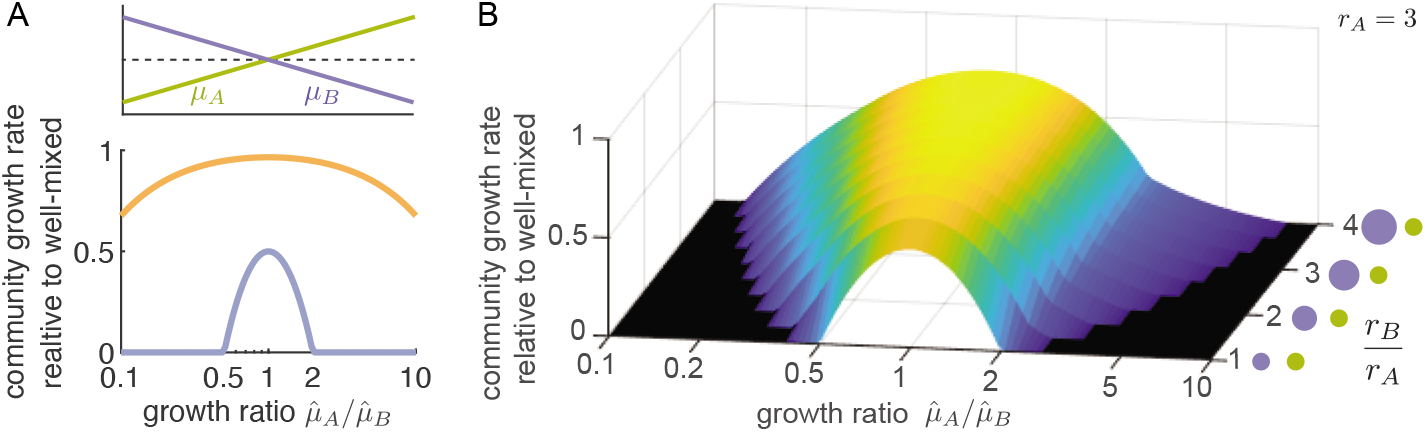
Small interaction neighborhoods can lead to community collapse. (A) When the neighborhood size is small (*r_A_* = *r_B_* = 3, purple line), asymmetric communities, where the cell types have different maximum growth rates, grow poorly. When the asymmetry is too large, communities cannot grow in a spatially structured environment even though they could grow in well-mixed environments. When the neighborhood size is large (*r_A_* = *r_B_* = 30, orange line), the growth rate of the spatial community is close to that of the well-mixed community even when there is an asymmetry. (B) Increasing the neighborhood size of just one the types is enough to increase the growth rate of spatial communities and prevent collapse. *r_A_* is held constant at 3, while *r_B_* is increased.

### Small interaction neighborhoods can lead to collapse of cross-feeding communities

Small interaction neighborhoods reduce the growth rate of cross-feeding communities. Here we show that this growth reduction is more pronounced when the two cell types have different maximum growth rates. We call such communities *asymmetric*. Our model predicts that the more asymmetric a cross-feeding community is, the more the community’s growth is hindered by small neighborhood sizes, to the point that the community collapses when neighborhood sizes are very small (Fig 5A). Increasing the neighborhood size of either one (Fig 5B), or both cell types (Fig 5A), can prevent the collapse of the community. This finding shows that there is a limit to the stability of cross-feeding interactions in spatially structured communities: cross-feeding cell types with different maximum growth rates that can stably coexist in well-mixed environments might not be able to survive in spatially structured environments. For cross-feeding communities, spatial structure can thus have a detrimental effect.

**Fig 5:**
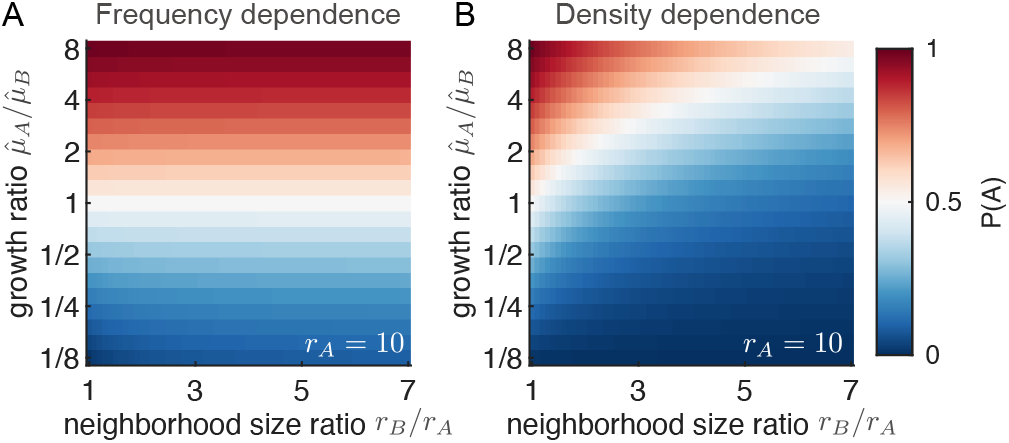
The neighborhood size strongly affects the equilibrium frequency when the growth rate of a cell depends on the absolute number of partner cells. (A) When the growth rate depends on the *frequency* of the partner type within the interaction neighborhood (frequency dependence), the equilibrium frequency of type A is almost completely determined by the ratio of the maximum growth rates (Eq. 1). (B) When the growth rate depends on the *number of cells* of the partner type within the interaction neighborhood (density dependence), the equilibrium frequency of type A depends both on the ratio of the maximum growth rates and on the ratio of the neighborhood sizes (SI Eq. 55). The slow growing type can still dominate the community when it has a much larger interaction range (blue region, top right). In both panels *r_A_* = 10.

### Small interaction neighborhoods affect other types of communities

So far, we have assumed that uptake and leakage rates differ between chemical compounds, but not between cell types. This assumption typically holds for microbial communities that consist of related strains and species, or for different cell types in a multicellular organism. In these cases all cell types likely share the same uptake and leakage pathways. However, uptake and leakage rates could differ both between chemical compounds and between cell types, e.g. because cell types use different uptake transporters or differ in their membrane permeabilities. Also in this general scenario we can derive expressions for the local rules, i.e. maximum growth rate and neighborhood size, as function of the molecular parameters of the molecular exchange (see Methods Eq. 7 and 8). With these local rules, we can derive analytical predictions for the community-level dynamics of any two species cross-feeding community.

So far we have focussed on cross-feeding systems were the growth rate of each cell type increases with the frequency of the other type within the interaction neighborhood. However, the growth rate of cells could depend on the partner type differently a nd o ur f ramework c an also be applied to these systems. We will illustrate this using two examples of cells interacting by non cross-feeding interactions. First, we consider a system where a type’s growth rate increases with the number, rather than the frequency, of partner type within the interaction neighborhood Fig 5). This is the case when cells exchange growth-affecting m olecules t hat a re n ot t aken up by the cells, but that are only sensed. If such molecules degrade, then the interaction range between cells can still be short even in the absence of uptake. Such molecules could for example be certain growth factors, which are rapidly inactivated by enzymes in the extracellular matrix that surround cells [37]. In this case, the type with the highest product of 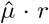 reaches the highest global frequency (see Supporting information). Therefore, the global composition of the community depends strongly both on the maximum growth rate and on the neighborhood size, rather than depending only on the maximum growth rate (Fig 5).

Second, we consider systems where two cell types inhibit each other’s growth. In such sytems, a cell’s growth rate decreases (linearly) with the frequency of the other type within the interaction neighborhood. We find that cells that inhibit each other grow slower when they are well-mixed compared to when they interact within a small neighborhood (see Supporting information). This happens because cells typically are surrounded by their own type, and thus have reduced interactions with the growth inhibiting cells.

In general our framework can be applied to any system of two cell types that affect each others growth by interacting within a finite range. As long as the growth rate of a cell depends linearly on the composition of its neighborhood, we can find an analytical solution for the steady state community properties; for non-linear functions, the model can be solved numerically (see Supporting information). The global composition of the community at steady state (*P* (*A*)) depends strongly on the chosen growth functions (Supporting information Eq. 47). However, the relation between the local and global composition of the community is always the same: i.e. for all growth functions we find that equation 3 holds (see Supporting information).

### Validating model using an experimental microbial community

We validate our mathematical framework using experiments with a bacterial cross-feeding com3 munity. This community consists of two cell types: one type unable to produce proline (Δ*P*), and one type unable to produce tryptophan (Δ*T*). We can parameterize our model in two ways: we can use the biophysical parameters describing the amino acid exchange (arrow 3 in Fig 1, Table S2) to calculate the neighborhood size (*r*, Eq. 4), and the maximum growth rate of a cell 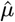 Eq. 6). Or, alternatively, we can parametrize the model using the measured interaction range and measured maximum growth rate (arrow 2 in Fig 1). The predictions obtained from the two alternative parameterization match closely (Table S3).

Our model recovers the experimentally observed community-level dynamics. In our experiments, the frequency of the Δ*T* type converges over time to a stable frequency across 22 communities (Fig 6A). Our model predicts similar dynamics: both pair approximation (Fig 6B) and simulations (Fig 6C) show that the community reaches a stable equilibrium. From Eq. 1 we find a predicted equilibrium frequency of Δ*T* of 0.20, which matches well with both the experimentally observed value of 0.19 and cellular automaton simulations value of 0.22 (Fig 6D). Therefore our model can quantitatively predict the global fraction of cell types based on biochemical parameters.

**Fig 6:**
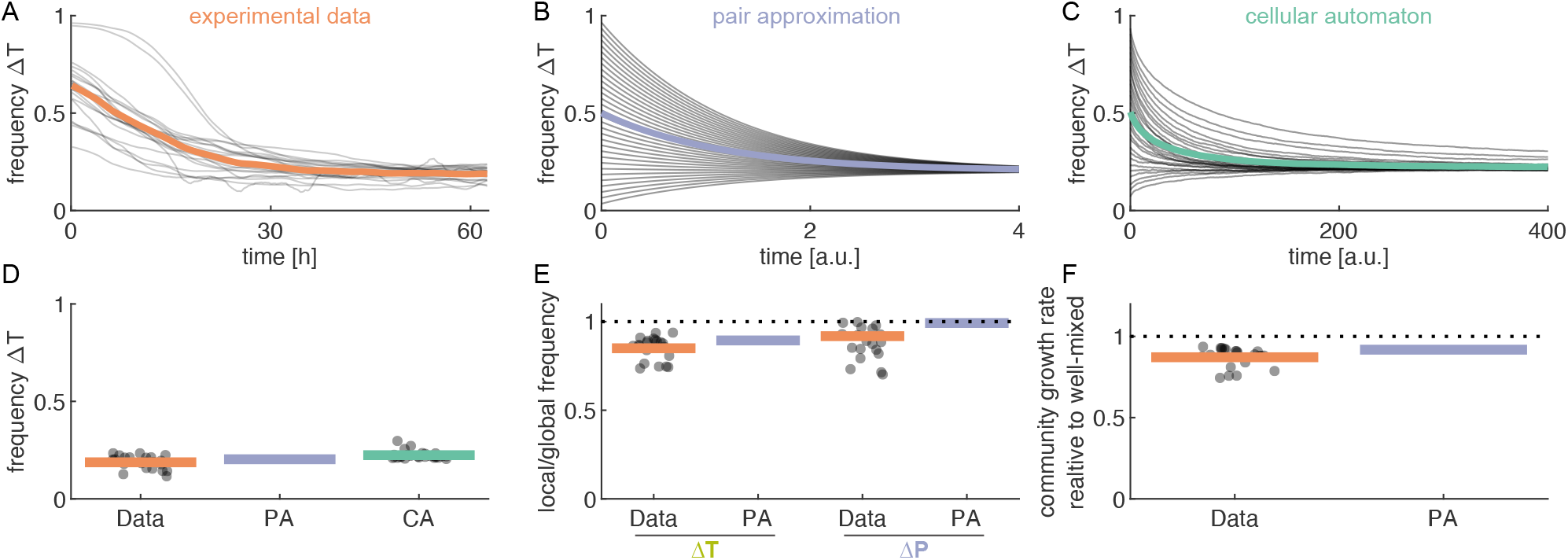
Our mathematical framework can explain experimentally observed community properties. The model parameters for the experimental community were calculated from the biophysical model (Table S2). (A) Experimental communities approach a stable frequency of Δ*T*. Individual communities (thin lines, n=22) and their average value (thick line) are shown. The initial frequency of Δ*T* in the communities is determined by the initial number of cells that enter the microfluidic growth chambers and the subsequent growth before the start of image acquisition; as a result it cannot be controlled experimentally. (B) Pair approximation predicts a unique stable equilibrium. The dynamical equations (Supporting information Eq. 34-36) were solved numerically starting from different initial frequencies. Time is in arbitrary units (a.u.). (C) Cellular automaton simulations also reach a unique stable equilibrium. (D) The observed equilibrium frequency of Δ*T* is consistent with the model predictions. Data: *P* (*A*)=0.19 (95% confidence interval (CI): 0.17–0.20), Pair-approximation: *P* (*A*)=0.20, and cellular automaton: *P* (*A*)=0.22 (CI: 0.22–0.23). The difference between the model prediction and data is less than 8%. (E) The frequency of the partner cell type within the interaction neighborhood (local frequency) is lower that the global frequency because cells are surrounded by their own offspring. Dots show measurements for 21 replicate communities, bar shows mean value. Pair approximation predicts (Eq. 3) a decrease in frequency of 0.99 for Δ*P* and 0.89 for Δ*T*, the experimental values are 0.87 (CI: 0.85–0.90) and 0.85 (CI: 0.82–0.88), respectively. The difference between the model prediction and data is less than 13%. (F) The average growth rate of the community is reduced due to cell clustering. An in-silico (see Methods) shows that the growth rate in clustered communities, with experimentally observed spatial arrangements, is reduced by a factor of 0.87 (CI: 0.85–0.90) compared to randomized communities, where cell clusters were disrupted (Data reproduced from [11]). Pair approximation predicts a decrease by a factor of 0.92 (see Supporting information Table S3); the difference between the model prediction and data is less than 6%.

Our model can also qualitatively capture the local dynamics observed in the experimental microbial communities. Cells tend to be surrounded by their own type, thus reducing the local partner frequency (Eq. 3). This is also observed in our experimental data: for both Δ*P* and Δ*T*, the local frequency of the partner type is reduced compared to the global frequency (Fig 6E). As a result of this decrease in the partner frequency, our model predicts that spatially structured cross-feeding communities, where cells become surrounded by their own type, have a lower average growth rate than well-mixed communities (Supporting information Eq. 52-53). Experimentally we cannot directly compare the growth in spatially structured communities with those in well-mixed communities. However, we previously used an individual-based biophysical model to perform an equivalent in-silico experiment: we compared the predicted growth rate for *clustered communities*, in which the spatial arrangement of cells was based on experimental measurements, with *randomized communities*, in which the spatial arrangement of cells was randomized [11]. We found that the growth rate in the real, clustered, communities was reduced compared to the randomized communities, in agreement with the prediction of the pair approximation (Fig 6F).

Quantitatively, our predictions for the local dynamics (Fig 6E & F) are not as accurate as those for the global dynamics (Fig 6D). This is likely due to the simplified representation of cells and space in the model, which is needed for the sake of analytical tractability. However, our model quantitatively predicts the global dynamics and captures the local dynamics. This means the model offers a helpful abstraction to reduce multiple biochemical parameters into few key combinations.

## Discussion

We developed a mathematical framework that can make quantitative predictions for the dynamics of spatially structured communities consisting of two interacting cell types (Fig 1). We primarily focussed on microbial cross-feeding systems and used pair approximation to derive analytical predictions for the community-level properties from knowledge of the local interaction rules (Fig 2). These rules are defined by two fundamental quantities: the size of the interaction neighborhood and the maximum growth rate that cells achieve when they are completely surrounded by the partner type; both quantities typically differ for the two cell types in the community.

We showed that these local interaction rules can be directly derived from key biophysical parameters of the underlying molecular mechanisms, such as uptake and leakage of the exchanged molecules (Fig 1). We worked out expression for the local interaction rules from the biophysical parameters for a number of scenarios. We validated this framework using an experimental community of two cross-feeding bacteria and showed that our model reproduces the experimentally observed community-level properties (Fig 6). Our framework elucidated how the local interaction rules arise from the molecular exchanges, and how the local interaction rules scale up to determine community properties. Generally, the framework we developed allows to scale up from molecular-level properties, to individual-level properties, to community-level properties.

For spatial cross-feeding communities, we found that the local and global properties can change independently. The global composition of the community (i.e. the equilibrium frequency of the two types) is set by the ratio of the maximum growth rates, which mostly depends on the leakage rates of the exchanged metabolites. In contrast, the composition of the local neighborhood, the one that matters the most for the cells, is set by the neighborhood size, which mostly depends on the ratio of the uptake and diffusion rates. Different biophysical parameters thus control the global and the local properties of the community.

Small neighborhood sizes reduce the frequency of the partner type around each cell. This happens because cells place their offspring close by in space. In our model we assumed that cells cannot actively move. However, cells could overcome the negative effect of having a small neighborhood size if they could actively move to locations with a higher frequency of the partner type. Without such active movement, cross-feeding cells in a spatial system grow slower than in equivalent well-mixed system (Fig 4). This reduction in growth rate can even lead to the collapse of the community: cells in asymmetric communities (i.e. communities where the two type have different maximum growth rates) can stably coexist in well-mixed system, yet they might not coexist in a spatial system, if they interact at small ranges (Fig 4). Moreover, dimensionality matters: given a fixed interaction range (*R*, determined by a set of molecular parameters) cells growing in two dimensional colonies typically have fewer neighbors (*r* α *R*^2^) compared to cells growing in three dimensional structures (*r* α *R*^3^). Our framework thus predicts that communities will grow slower in two-dimensional colonies than in three-dimensional clusters. Our work joins a body of work showing that the spatial dimensionality of a system has an influence on cellular interactions [25].

Previous work suggested that short-range interactions provide an advantage to cooperative interactions, because they separate cooperative types from non-cooperators [29, 38–40]. In general, when it is good to be surrounded by your own type, we expect short-range interactions to be beneficial, as for example when two cell types inhibit each other. When it is good to be surrounded by the other type, we expect short-range interactions to be detrimental, as for example when two cell types exchange beneficial resources. When a community consists of more than two cell types the situation can be more complex. Previous studies have shown that short-range interaction can be beneficial for cross-feeding communities that contain non-producing cells: the short-range interactions harm the producing cells, but they harm the non-producers even more, and thus prevent them from taking over [22, 41, 42].

Our framework can also model systems beyond the bacterial world, as long as they are composed of two interacting types. For example, a central question in tissues homeostasis is how two (or more) different cells types can control each other’s growth by exchanging diffusible growth factors to maintain proper tissue functioning [43, 44]. These cellular system are typically spatial, in the sense that interactions act on a finite distance. From the point of view of modeling, spatial structure introduces complexity in the mathematical representations, as well as increasing the parameter space of models. Our framework provides a simple approach to study the equilibrium properties and can assist in the design of synthetic systems and tissue engineering, as it allows for the prediction of system-level properties from molecular scale parameters [45].

Overall, our model offers a versatile representation of spatial system of interacting cells that can be adapted to various types of interactions. Many biological systems are spatially structured and multi-scale: they consist of individual entities (e.g. cell types or species) that interact with each other in space. Interactions at different levels determine the global properties of the system (e.g. multicellular organism or microbial community). To understand such biological systems, it is important to scale between levels of organization. Our work provides a contribution to this effort by creating a mathematical framework that can scale from molecular mechanisms, to local interaction rules, to global system properties.

## Methods

### Model assumptions

- Two cell types, A and B, fully occupy a regular directed graph. Type A interacts with *r_A_* neighbors, type B interact with *r_B_* neighbors, where *r_B_* ≥ *r_A_*.
- A cell’s growth rate increases linearly with the frequency of the other type within its interaction neighborhood.
- Individuals reproduce with a probability proportional to their growth rate, and their off-spring replaces a random neighbor within the replication neighborhood.
- The replication neighborhood is identical to the smallest interaction neighborhood *r_R_ = r_A_*.

We used pair approximation to derive analytical predictions and a cellular automaton to run simulations of the model.

### Pair approximation

We track the number of all pairwise links *N_X←Y_*, where *X, Y* ∈ *A, B*. The total number of links changes through time, because the neighborhood sizes are different for the two cell types (see Supporting information). This is in contrast to previous work were the two neighborhood sizes are the same and the total number of links remains constant. Two events can change the number of links: a type A cell reproduces and replaces a B neighbor, with rate *T* ^+^, or a type B cell reproduces and replaces an A neighbor, with rate *T* ^−^. During a *T* ^+^ event the number of A cells thus increases by one, and during a *T ^−^* event it decreases by one. The rate *T* ^+^ is given by:

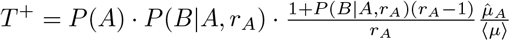

where the first factor gives the probability of choosing a type A focal cell, the second the probability that this A cells has a type B neighbor, and the third the average growth rate of a type A cell that has at least 1 type B neighbor, relative to the average population growth rate ⟨*μ*⟩, which is given by:

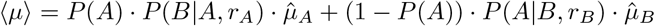

Similarly, for *T ^−^* we have:

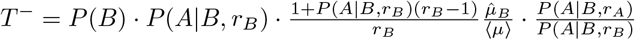

The extra factor at the end is the probability that a type A neighbor that is in the (larger) interaction neighborhood is also part of the smaller replication neighborhood. We can express all probabilities as function of the number of links:

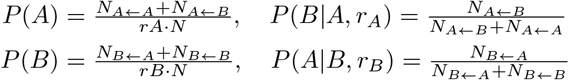

where *N* is the total number of cells in the systems.

When a *T* ^+^ or *T* ^−^ event happens, the number of *X* ← *Y* link changes by 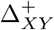 and 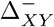, respectively. These changes can be fully expressed as function of *N_X←Y_*, as we show in the Supporting information. We can then write the dynamical equation for *N_X←Y_* as:

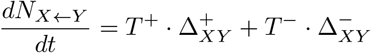

We can solve these equation numerically to obtain the temporal dynamics, and we can solve them analytically for the steady state, see Supporting information for full derivation.

### Model parametrization

To parametrize the model, we need to specify the size of the interaction neighborhood *r* and the maximum growth rate of a cell 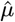. We previously showed that that interaction-range (*R*) of a cell (the distance over which amino-acids can be exchanged) is given by [11]:

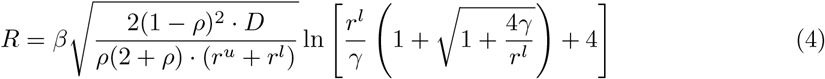

where *β* and *γ* are constants, *ρ* is the volume fraction occupied by cells, and *D*, *r^u^*, and *r^l^* are the rates of diffusion, uptake, and leakage of the exchanged metabolite (see Supporting information for details and parameter values). The number of neighbors in 2D can be directly calculated from the interaction range:

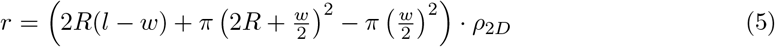

where *l* and *w* are the average length and width of cells, and *ρ*_2*D*_ is the number of cells per area. A similar expression can be derived for the number of neighbors in a three-dimensional system (see Supporting information Eq. 22).

We previously showed that that maximum growth rate of a cell 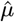 is given by [11]:

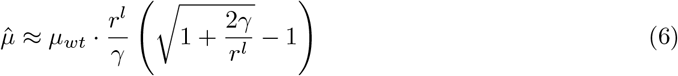

### Generalized model parameterization

In the previous section we assumed that the uptake and leakage rates differ between molecules, but not between cell types. Here we relax this assumption and show the local interaction rules for systems where rates differ both between molecules and cell types. In the Supporting information we show that in this case the maximum growth rate is given by:

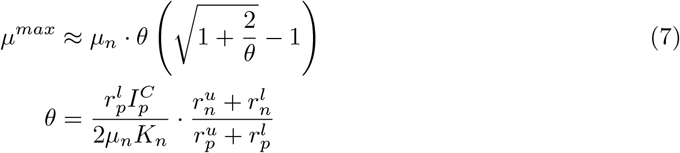

Here the subscript *n* refers to parameters of non-producing cells, while *p* refers to parameters of the producing cells. 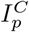 is the internal concentration of the produced molecule inside prodcuer cells, *K_n_* is the Monod constant of the consumer cells and *μ_n_* is the highest rate at which consumer cells can grow when the exchanged molecule is provided in excess. The first term in the constant *θ* measures the leakage flux in producing cells 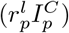 relative to the flux needed by non-producing cells to grow well (2*μ_n_K_n_*). The second term corresponds to the effective uptake rate (active transport with rate 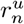 together with diffusion across the membrane with rate 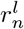) in non-producing cells relative to that in producing cells. In the case where cell types have identical rates, 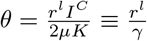 and we thus recover equation 6.

Moreover, we can show (see Supporting information), that the growth range is given by:

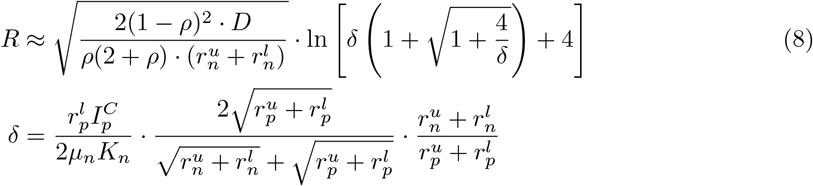

The first term in the constant *δ* measures the leakage flux in producing cells relative to the flux needed by non-producing cells to grow well. The second term corresponds to the diffusion length scale 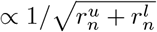 in regions occupied by non-producing cells, relative to the average diffusion length scale. The third term corresponds to the effective uptake rate in non-producing cells relative to that in producing cells. In the case where cell types have identical rates, 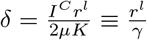 and we thus recover equation 4.

### Cellular automaton

In the cellular automaton cells are placed on a square grid of size 100 × 100 with periodic boundary conditions. Cells interact within an extended Moore neighborhood with range *d_X_*, and thus have a neighborhood size of (2*d_X_* + 1)^2^ − 1. For each cell we calculate the growth rate (using linear growth function) and we randomly pick a cell to reproduce with a probability that is proportional to its growth rate. We then randomly replace one of its neighbors. The grid was initialized with a random arrangement with a given frequency of the two types. We parameterized the experimental community by choosing *d*_Δ*T*_ = 1 (8 neighbors) and *d*_Δ*P*_ = 5 (120 neighbors). Simulations are implemented in C++.

### Experimental communities

The experimental methods are described in detail in reference [11], here we summarize the most relevant details. The community consists of two strains of *Escherichia coli*: ΔT: MG1655 trpC::frt, PR-sfGFP and ΔP: MG1655 proC::frt, PR-mCherry. The deletion of *trpC* and *proC* prevents the production of tryptophan and proline, respectively. Cells are labeled with constitutively expressed fluorescent proteins.

Cells were grown in microfluidic chambers of 60×60×0.76*μ*m, the small height forces cells to grow in a monolayer. The chambers open on one side into a feeding channel of 22*μ*m high and 100*μ*m wide. The microfluidic devices were fabricated from Polydimethylsiloxane (PDMS) using SU8 photoresist molds. Overnight cultures of the two strains (started from single colonies) were mixed in a 1:1 volume ratio and loaded into the microfluidic devices by pipette. Cells were grown on M9 medium (47.8mM Na_2_HPO_4_, 22.0 mM KH_2_PO_4_, 8.6 mM NaCl and 18.7 mM NH_4_Cl) supplemented with 1 mM MgSO_4_, 0.1 mM CaCl_2_, 0.2% glucose, and 0.1% Tween-20. For the first 10h the medium was supplemented with 434 mM of L-proline and 98 mM of L-tryptophan to allow cells to grow independently and fill the chambers. Subsequently, cells were grown without externally supplied amino acids.

The growth of the communities was followed using time-lapse microscopy. Phase contrast and fluorescent images were taken every 10min using fully automated Olympus IX81 inverted microscopes, equipped with a 100x NA1.3 oil objective, a 1.6x auxiliary magnification, and a Hamamatsu ORCA flash 4.0 v2 sCMOS camera. The sample was maintained at 37°C with a microscope incubator.

### Image analysis

Microscope images were analyzed using Vanellus (version v1.0 [46]). Images were registered, cropped to the area of the growth chambers, and deconvoluted. Images were segmented on the fluorescent channels using custom build Matlab routines [11] or on a combination of the phase contrast and fluorescent channels using Ilastik (version v1.3.3 [47]). Cell tracking was done using custom build Matlab routines [11] followed by manual correction. Cell length and width were measured by fitting an ellipse to the cell shape and taking the major and minor axis length, respectively. Cell growth rates were estimated by fitting a linear regression to the log transformed cell lengths over a 40min time window [11].

### Data analysis

For each chamber we estimated *P* (Δ*T*) as the number of pixels occupied by Δ*T* cells divided by the total number of pixels occupied by cells. We estimated *P* (Δ*T* |Δ*P, r*_Δ*P*_) by first calculating the local Δ*T* frequency for each Δ*P* cell, and averaging this over all Δ*P* cells within the chamber. The local frequency was calculated as the number of pixels occupied by Δ*T* cells divided by the total number of pixels occupied by cells, considering only pixels within 12.1*μ*m (the interaction range of Δ*P*) of the cell perimeter. *P* (Δ*P* |Δ*T, r*_Δ*T*_) was estimated in similar way, using the interaction range of 3.2*μ*m for ΔT.

The difference between the average growth rates of *clustered* and *randomized communities* was estimated using an in-silico experiment. For the clustered communities experimentally observed spatial arrangements of 22 communities were converted to a 40×40 square grid. Each grid site was assigned the cell type that occupied the majority of the corresponding pixels in the real image. Grid sites that remained empty after this procedure were randomly assigned one of the types. This grid was used as input for a previously described individual based model that can predict single cell growth rates [11]. The resulting growth rates were averaged over all cells to obtain the average growth rate for the clustered communities. Subsequently the 40×40 grids were randomized, keeping the frequency of the two cell types constant but permuting their locations. We calculated the average community growth rate for 20 independent randomizations, and we assigned the average over all these permutations as the growth rate of the randomized communities.

## Code and Data Availability

The Matlab code used to solve the dynamical equations and Mathematica worksheet used the derive the analytical expressions are available on Github: https://github.com/simonvanvliet/Pair_Approximation_Community_Dynamics.git; the data is available at the ETH repository: https://doi.org/10.3929/ethz-b-000368486.

## Author contributions

SvV: Conceptualization, Methodology, Formal analysis, Writing – original draft. CH: Methodology, Formal analysis, Writing – review & editing. MA: Conceptualization, Writing – review & editing. ADC: Conceptualization, Methodology, Formal analysis, Writing – original draft. The authors declare no conflicts of interest.

## Acknowledgements

We thank Michael Doebeli and the members of the Microbial Systems Ecology group, the Doebeli lab, and the Hauert lab for valuable input and discussion. SVV was funded by the SNSF Postdoc Mobility fellowship nr. 175123, CH was funded by the Natural Sciences and Engineering Research Council of Canada (NSERC) grant nr. RGPIN-2015-05795, MA and ADC were funded by SNSF grant nr. 31003A_169978, Eawag, and ETH Zurich, and ADC was funded by the Office of Naval Research grant nr. ONR N00014-17-1-3029.

## Notes

### Competing Interest Statement

The authors have declared no competing interest.

### Summary of Updates

Revision of main-text, extended biophysical model to 3D and asymmetric communities.

