## Supplemental Information for "Global dynamics of microbial communities emerge from local interaction rules"

### Supporting Information for: Global dynamics of microbial communities emerge from local interaction rules

4 Supplementary Discussion 23

5 Supplementary Tables 25

6 References 27

### 1 Local interaction rules for cross-feeding communities

We previously showed that the local interaction rules for cross-feeding communities can be fully specified with two quantities: the range over which cells can obtain nutrients, i.e. the interaction range $R$, and the maximum rate at which cells grow when they are fully surrounded by the partner type, $\hat{\mu}$ (1). Using a biophysical model, we showed that these two quantities can be derived from the biophysical parameters underlying the exchange of metabolites. We previously derived these results assuming that communities grow in two-dimensional structures and consist of closely related cell types that share the same pathways for the uptake and release of nutrients. Here we will generalize our biophysical model by extending it to three-dimensional communities, and to communities consisting of cell types that can differ substantially in their pathways for the uptake and release of nutrients. The extended model presented here is thus generally applicable to any cross-feeding community consisting of two cell types.

#### 1.1 Summary of previous results for two-dimensional communities

Here, we will briefly summarize the local rules for two-dimensional cross-feeding communities which we derived previously, for the full derivation we refer the reader to the supplementary information of reference (1).

##### 1.1.1 Interaction range

The first essential parameter that describes the local rules of cross-feeding communities is the interaction range $R$, which specifies the distance, measured from the cell surface, over which amino-acids are primarily exchanged. We previously derived an analytical approximation for this interaction range based on the molecular parameters of the underlying nutrient exchange (1). Specifically, for simple spatial arrangements where the two cell types are separated by a straight interface, we could calculate how the growth rate decreases away from the interface between the two cell types. We calculated the distance over which the growth rate decreases by 50% (i.e. the growth range) and showed that this distance is directly proportional to the interaction range; the interaction range is a measure of the distance over which amino acids can be exchanged in an arbitrarily complex spatial arrangement (1). We found that the interaction range can be approximated as:

$$R = \beta \sqrt{\frac{2(1-\rho)^2 \cdot D}{\rho(2+\rho) \cdot (r^u + r^l)}} \ln \left[ \frac{r^l}{\gamma} \left( 1 + \sqrt{1 + \frac{4\gamma}{r^l}} \right) + 4 \right] \tag{1}$$

where $\beta$ and $\gamma$ are constants, $\rho$ is the 3D cell density (i.e. the volume fraction occupied by cells), and $D$, $r^u$, and $r^l$ are the rates of diffusion, uptake, and leakage of the exchanged metabolite. The square root term can be interpreted as the distance that a molecule travels before it is taken up by a cell, and this depends primarily on the ratio of diffusion rate and uptake rate and the cell density. The constant $\beta$ is the constant of proportionality between the growth range and the interaction range, and we previously showed that $\beta \approx 0.88$.

The constant $\gamma = 2\mu^{wt}/\mathcal{I}^C$ is species specific, but does not depend on the properties of the exchanged molecules. Here, $\mu_{wt}$ is the maximum growth rate of a wild type cell that can produce all essential metabolites. $\mathcal{I}^C = I^C/K_M$ is the internal concentration of the essential metabolite in a producing cell ($I^C$) relative to the Monod constant of the growth curve ($K_M$; growth is assumed to follow Monod kinetics as function of the internal concentration ($I$) of the essential metabolite: $\mu(I) = \frac{\mu_{wt} I}{I + K_M}$). The factor $\frac{r^l}{\gamma} = \frac{r^l I^C}{2\mu^{wt} K_M}$ can be interpreted as the flux of metabolites leaked into the environment, relative to the flux of metabolites used for growth. It is thus a measure of leakiness: if it is close to 1, a large fraction of the essential metabolite is lost to the environment, while if it is much smaller than 1 most of the essential metabolite is kept within the cell and used for growth. To derive Eq. 1 we assumed that $r^l \ll \mu_{wt}$ and $I^C \gg K_M$. These assumptions thus state that cells have limited leakiness. We previously showed that these assumptions are compatible with the measurements from an experimental cross-feeding community (1).

###### 1.1.2 Maximum growth rate

The second essential parameter that describes the local rules of cross-feeding communities is the maximum growth rate that cells can obtain when they are fully surrounded by the other cell type, $\hat{\mu}$. We previously showed that (1):

$$\hat{\mu} \approx \mu_{wt} \cdot \frac{r^l}{\gamma} \left( \sqrt{1 + \frac{2\gamma}{r^l}} - 1 \right) \tag{2}$$

##### 1.2 Local interaction rules for three-dimensional communities

The analytical expressions for the local interaction rules described above were originally derived for two-dimensional systems, however we will show here that they also hold for three-dimensional systems. Specifically, the derivation for the maximum growth rate (Eq. 2) only assumed that isolated non-producing cells, fully surrounded by producing cells do not substantially change the external concentration of the exchanged compound; this assumption holds equally in 2D and 3D. The analytical expression for the growth range (Eq. 1) also generalizes to 3D. The primary assumption we made to derive this result is that the interface between the two cell types is quasi-1D; in 2D systems this assumption means the two types are separated by a straight line, and in 3D it means the two types are separated by a plane. For these very simple configurations we can then calculate the growth range using the same quasi-1D approximation. Even though the growth range $R$ is identical for 2D and 3D systems, the neighborhood size $r$ does depend on dimensionality. For spherical cells, the number of cells within a distance $R$ of a central cell is proportional to $R^2$ in 2D and proportional to $R^3$ in 3D. For rod-like cells the relation between the number of neighbors $r$ and the growth range $R$ is more complicated, as we will show below (Eq. 21 for 2D and 22 for 3D), however it is still true that the number of neighbors, at a constant growth range, is higher in 3D than in 2D systems.

For more complex spatial arrangements we cannot calculate the growth range analytically, however we can use an individual based model that we developed previously to numerically calculate the interaction range for any arbitrary 2D spatial arrangement (1). Here we extended this individual based model to 3D. This is done by replacing the square 2D grid with a cubic 3D grid, and by using a 3D instead of 2D Laplacian to calculate the diffusion of metabolites (see Eq. 15 and 16 in the SI of reference (1)). We used this model to calculate the interaction range for 2D and 3D communities with varying spatial arrangements. The grid is initialized by randomly assigning each grid point to one of the two cell types. To vary the patch size, we pick a random cell and let it replace one of its randomly chosen neighbors; for each *replication cycle* this is done until all cells on the grid have replaced one of their neighbors. The procedure can be repeated for more cycles to achieve larger patches on average (see Fig S1A). For each grid we then calculate the steady state growth rates of all cells, and then use these growth rates to calculate the interaction range using the correlation method we previously developed. In short: the growth rate of a cell is correlated with the frequency of the partner type within a given radius. The interaction range is defined as the radius at which this correlation is maximal (see reference (1) for details).

Using the individual based model we can show that the interaction range we derived is very similar for 2D and 3D systems. Fig S1B shows the calculated interaction range for 2D and 3D arrangements of a symmetric community for varying patch sizes. The estimated interaction ranges are very similar between 2D and 3D systems, and both are close to the analytically calculated growth range (Eq. 1). This thus shows that our estimate of the range over which cell can interact holds for both 2D and 3D systems.

#### 1.3 Local interaction rules for communities consisting of dissimilar cell types

Previously we assumed that uptake and leakage rates differ between chemical compounds, but not between cell types (1). This assumption holds whenever all cell types share the same pathway for uptake and leakage, as it is the case for microbial communities that consist of closely related strains or for different cell types in a multicellular organism. However, in general uptake and leakage rates could differ both between chemical compounds and between cell types, due to difference in uptake and leakage pathways. Here we will derive the local rules, i.e. the maximum

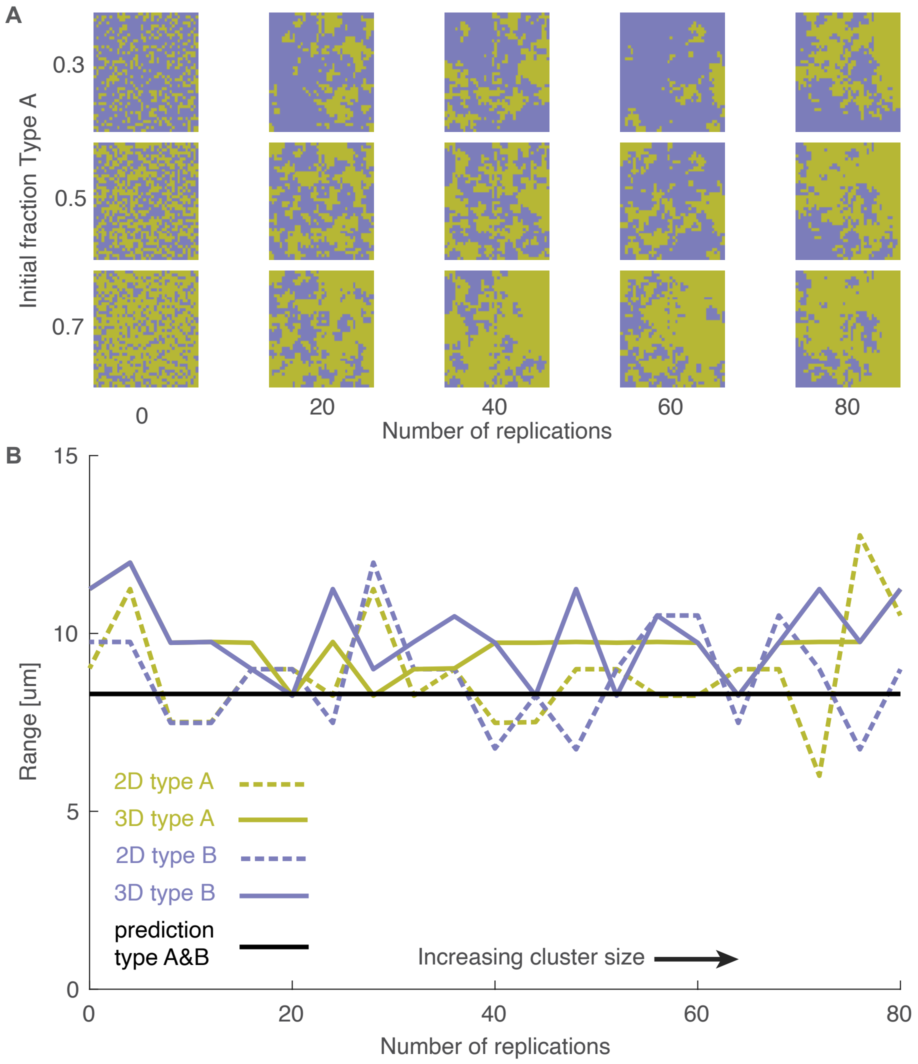

**Fig. S1:** The interaction range is similar for 2D and 3D communities and closely matches the analytical prediction for the growth range. The interaction range was calculated based on in-silico data for a community consisting of two cell types (A and B) with identical molecular rates. To calculate the interaction range, we used a previously developed individual based model (1) to predict the growth rate of all cells growing either in a 2D or 3D community; then used a correlation method (1) to estimate the interaction range for both cell types. The interaction range is compared to the growth range, which is calculated analytically from Eq. 1. For each parameter set, we simulated 100 communities of 40x40 cells (2D) or 40x40x40 cells (3D), with closed (no-flux) boundaries on all sides as described previously (1). The grids were seeded by randomly assigning the two cell types with a frequency of type A between 0.2 and 0.8. To implement larger patches we allowed all cells (in random order) to replace one of their neighbors (chosen at random). In total we performed 0 to 80 of these replication cycles. The more replication cycles the larger the average patch size. A) Typical spatial arrangements as function of the initial frequency of type A (y-axis) and the number of replication cycles (x-axis), the central slice of the 3D grid is shown. B) The calculated interaction range is shown as function of the number of replication cycles (i.e., patch size) and compared to the analytical prediction for the growth range. Parameters: $r_A^u = r_B^u = 3.58\ 1/s$, $r_A^l = r_B^l = 3.58 \cdot 10^{-6}\ 1/s$, $D_A = D_B = 7.17 \cdot 10^2\ \mu m^2/s$, all other parameters as in table S1.

growth range and interaction range, for such systems. We closely follow reference (1) to derive these quantities and we refer the reader to that document for more details on the derivation.

We track the internal $I$ and external $E$ concentration of the exchanged compounds. For notational simplicity, we focus on the exchange of a single compound and we write down the equations for the internal $I_p$ and external $E_p$ concentration for cells that can produce this compound:

$$\frac{\partial I_p}{\partial t} = 0 \tag{3}$$

$$\frac{\partial E_p}{\partial t} = -\alpha \cdot r_p^u \cdot E_p + \alpha \cdot r_p^l \cdot (I_p - E_p) + D^{eff} \nabla^2 E_p \tag{4}$$

where $r_p^u$ and $r_p^l$ are the uptake and leakage rates of the exchanged compound of the producing cells, respectively, and where $D^{eff} = \frac{(1-\rho)\cdot D}{(1+\frac{\rho}{2})}$ is the effective diffusion constant in a crowded environment, and where $\alpha = \frac{V_{in}}{V_{out}} = \frac{\rho}{1-\rho}$ is the ratio between the intra- and extracellular volume. Similarly, we write the equations for the internal $I_n$ and external $E_n$ concentration for cells that cannot produce this compound:

$$\frac{\partial I_n}{\partial t} = r_n^u \cdot E_n - r_n^l \cdot (I_n - E_n) - \frac{\mu_n \cdot I_n}{K_n + I_n} \cdot I_n \tag{5}$$

$$\frac{\partial E_n}{\partial t} = -\alpha \cdot r_n^u \cdot E_n + \alpha \cdot r_n^l \cdot (I_n - E_n) + D^{eff} \nabla^2 E_n \tag{6}$$

where $r_n^u$ and $r_n^l$ are the uptake and leakage rates of the exchanged compound of the non-producing cells, respectively, $K_n$ is the Monod constant for the non-producing cells (the concentration at which cells can grow at half-maximum rate), and $\mu_n$ is the growth rate that auxotrophic cells can reach when the exchanged compound is non-limiting.

By solving Eq. 5 for the temporal steady state we can find an expression for the internal concentration as function of the external concentration inside consumer cells:

$$I_n(E_n) = \frac{(r_n^u + r_n^l)E_n - r_n^l K_n + \sqrt{((r_n^u + r_n^l)E_n + r_n^l K_n)^2 + 4(r_n^u + r_n^l)\mu_n K_n E_n}}{2(\mu_n + r_n^l)} \tag{7}$$

##### 1.3.1 Maximum growth rate

Here we derive the analytical expression for the growth rate of a non-producing cell surrounded by a large number of producing partners. We assume that the single non-producing cell has a negligible influence on the external concentration of the exchanged molecules; thus this concentration is identical to that in an area fully occupied by producers cells and can be found by solving Eq. 3 and 4 at steady state :

$$E^{max} = \frac{r_p^l}{r_p^u + r_p^l} \cdot I_p^C \tag{8}$$

where $I_p^C$ is the internal concentration of the produced compound inside the producer cell. We can then calculate the growth rate of the consumer cell using:

$$\mu^{max} = \frac{\mu_n I_n(E^{max})}{K_n + I_n(E^{max})} \tag{9}$$

where $I_n(E^{max})$ is found by substituting $E_n$ in Eq. 7 with $E^{max}$ as given by Eq. 8. Although an analytical expression can be obtained, it is rather complex and we therefore follow reference (1) and further simplify the expression by assuming:

**Assumption 1** $\frac{I_p^C}{K_n} \gg \frac{r_n^l}{r_p^l} \cdot \frac{r_p^u + r_p^l}{r_n^u + r_n^l}$. *This is identical to Assumption 1 in reference (1), with the additional requirement that the difference in uptake and leakage rates between the two cell types is not too large.*

**Assumption 2** $r^l \ll \mu^{aux}$. *This is identical to Assumption 2 in reference (1).*

Using these assumptions we find the following expression for the maximum growth that non-producing cells can obtain in a cross-feeding consortium:

$$\mu^{max} \approx \mu_n \cdot \theta \left( \sqrt{1 + \frac{2}{\theta}} - 1 \right) \tag{10}$$

$$\theta = \frac{r_p^l I_p^C}{2\mu_n K_n} \cdot \frac{r_n^u + r_n^l}{r_p^u + r_p^l} \tag{11}$$

The first term in the constant $\theta$ measures the leakage flux in producing cells ($r_p^l I_p^C$) relative to the flux needed by non-producing cells to grow well ($2\mu_n K_n$). The second term corresponds to the effective uptake rate (active transport with rate $r_n^u$ together with diffusion across the membrane with rate $r_n^l$) in non-producing cells relative to that in producing cells. In the case where cell types have identical rates, $\theta = \frac{r^l I^C}{2\mu K} = \frac{r^l}{\gamma}$ and we thus recover Eq. 2 (Eq. 25 in SI of reference (1)) which we previously derived for communities where both cell types have the same uptake and leakage rates. Eq. 10 increases monotonically with $\theta$ showing that the maximum growth rate of non-producing cells increases with the leakage rate of producer cells ($r_p^l$) and the uptake rate of non-producing cells ($r_n^u$), while it decreased with the uptake rate of producer cells ($r_p^u$).

##### 1.3.2 Growth range

Next we will re-derive the growth range for communities in which the cell types differ in their uptake and leakage rates. The derivation closely follows reference (1). We consider a scenario where the two cell types are symmetrically arranged i.e. when the two cell types are separated by a straight line in 2D or by a flat plane in 3D. This reduces the problem to one-dimension, $x$, which measures the distance of a cell to the interface. Producing cells are located at $x < 0$ and non-producing cells at $x > 0$. We will here derive an analytical approximation for the growth rate of the non-producing cells by setting Eq. 4 and 6 to steady state:

$$\frac{d^2E}{dx^2} = \begin{cases} \frac{1}{r_{p0}^2}\left(E - \frac{r_p^l}{r_p^l + r_p^u} I_p^C\right) & \text{if } x < 0 \\ \frac{1}{r_{n0}^2}\left(E - \frac{r_p^l}{r_p^l + r_p^u} I_n(E)\right) & \text{if } x > 0 \end{cases} \tag{12}$$

where

$$r_{p0} = \sqrt{\frac{D^{eff}}{\alpha(r_p^u + r_p^l)}} \qquad r_{n0} = \sqrt{\frac{D^{eff}}{\alpha(r_n^u + r_n^l)}}$$

For $x > 0$ the analytical solution of Eq. 12 cannot be found due to the non-linear term $I_n(E)$ (given by Eq. 7), however we previously showed that this term can be ignored close to the interface (1). We thus solve the following approximate ODE:

$$\frac{d^2E}{dx^2} = \begin{cases} \frac{1}{r_{p0}^2}\left(E - \frac{r_p^l}{r_p^l + r_p^u} I_p^C\right) & \text{if } x < 0 \\ \frac{1}{r_{n0}^2} E & \text{if } x > 0 \end{cases} \tag{13}$$

This equation can be solved analytically to find:

$$E(x) = \begin{cases} C_1 \cdot e^{x/r_0p} + \frac{r_p^l}{r_p^l + r_p^u} I_p^C & \text{if } x < 0 \\ C_2 \cdot e^{-x/r_0n} & \text{if } x > 0. \end{cases}$$

We can solve for $C_1$ and $C_2$ by imposing continuity of concentration and flux at the interface and find:

$$E(x) = \begin{cases} \frac{I_p^C r_p^l \left(r_{0n} - r_{0p} e^{\frac{x}{r_{0p}}} + r_{0p}\right)}{\left(r_p^l + r_p^u\right)(r_{0n} + r_{0p})} & \text{if } x < 0 \\ \frac{I_p^C r_p^l r_{0n} e^{-\frac{x}{r_{0n}}}}{\left(r_p^l + r_p^u\right)(r_{0n} + r_{0p})} & \text{if } x > 0 \end{cases} \tag{14}$$

We now calculate an analytical approximation for the *growth range* ($R$), which is the distance from the interface where cells have 50% of the growth rate they have at the interface:

$$\mu(x = R) = \frac{1}{2} \cdot \mu(x = 0) \tag{15}$$

as $\mu = \frac{\mu_n \cdot I_n}{K_n + I_n}$ it follows that

$$I_n\Big|_{x=R} = \frac{I_n\Big|_{x=0}}{2 + I_n\Big|_{x=0}} \tag{16}$$

where $I_n\Big|_x \equiv I_n(E(x))$. Substituting Eq. 7 for $I_N(E)$ and Eq. 14 for $E(x)$ we can thus solve Eq. 16 for the growth range:

$$R = r_{0n} \log \left[ \frac{\left(4r_n^l - \delta r_p^l\right)\left(\sqrt{2\delta r_p^l \mu_n + \left(\frac{\delta r_p^l}{2} + r_n^l\right)^2} + \frac{\delta r_p^l}{2} + r_n^l\right) - 4\delta r_p^l\left(\mu_n + r_n^l\right)}{2r_n^l\left(2r_n^l - \delta r_p^l\right) - \delta r_p^l \mu_n} \right] \tag{17}$$

where

$$\delta = \frac{r_p^l I_p^C}{2\mu_n K_n} \cdot \frac{2r_{0n}}{r_{0n} + r_{0p}} \cdot \frac{r_n^u + r_n^l}{r_p^u + r_p^l} \tag{18}$$

$$= \frac{r_p^l I_p^C}{2\mu_n K_n} \cdot \frac{2\sqrt{r_p^u + r_p^l}}{\sqrt{r_n^u + r_n^l} + \sqrt{r_p^u + r_p^l}} \cdot \frac{r_n^u + r_n^l}{r_p^u + r_p^l} \tag{19}$$

The first term in the constant $\delta$ measures the leakage flux in producing cells ($r_p^l I_p^C$) relative to the flux needed by non-producing cells to grow well ($2\mu_n K_n$). The second term corresponds to the diffusion length scale $r_{0n}$ in regions occupied by non-producing cells, relative to the average diffusion length scale $(r_{0p} + r_{0n})/2$. The third term corresponds to the effective uptake rate (active transport with rate $r_n^u$ together with diffusion across the membrane with rate $r_n^l$) in non-producing cells relative to that in producing cells.

Using assumption 2 and

**Assumption 3** $\frac{I_p^C}{K_n} \gg \frac{r_n^l}{r_p^l} \cdot \frac{r_{0n}+r_{0p}}{r_{0n}} \cdot \frac{r_p^u+r_p^l}{r_n^u+r_n^l}$. *This is identical to Assumption 1 in reference (1), with the additional requirement that the difference in uptake and leakage rates between the two cell types is not too large.*

this can be simplified to:

$$R \approx \sqrt{\frac{D^{eff}}{\alpha(r_n^u + r_n^l)}} \cdot \ln\left[\delta\left(1+\sqrt{1+\frac{4}{\delta}}\right)+4\right] \tag{20}$$

In communities where both cell types have the same uptake and leakage rates, $\delta = \frac{I^C r^l}{2K\mu} = \frac{r^l}{\gamma}$, recovering our previous results (i.e. Eq. 1). Eq. 20 is a monotonically decreasing function of $r_n^u$, so the higher the uptake rate of the consumer, the lower the growth range. When $r_n^u$ and $r_n^l$ are held constant, Eq. 20 is a monotonically increasing function of $\delta$.

#### 1.4 Estimating neighborhood size from interaction range

Above we derived the local rules for cross-feeding systems by calculating the maximum growth range and the interaction range of cells. The interaction range determines the interaction neighborhood of a cell by specifying the maximum distance, measured from surface of a cell, over which cells can obtain nutrients. However, in the pair-approximation framework we derive below (see section 2) the interaction neighborhood has to be specified as the number of cells that are contained within it. These two metrics, i.e. the interaction range, measured in units of distance, and the neighborhood size, measured as a number of cells, can be converted into each other using a simple geometrical calculation.

For two-dimensional systems, we can estimate the number of neighbors ($r$, measured in number of cells) from the interaction range ($R$, measured in units of length) using measured values of the cell geometry and the cell area density $\rho_{2D}$ (i.e. the number of cells per area). We assume that cells (as seen in a 2D projection) can be described as rectangles with semi-spherical end-caps, with total length $l$ and width $w$. The number of cells within the interaction neighborhood (i.e. within the area within distance $R$ from the cell-surface) is then given by:

$$r = \left(2R(l-w) + \pi\left(R+\frac{w}{2}\right)^2 - \pi\left(\frac{w}{2}\right)^2\right) \cdot \rho_{2D} \tag{21}$$

We can make a similar estimate for three-dimensional systems. We assume that cells can be described as cylinders with semi-spherical end-caps, with total length $l$ and width $w$. The number of cells within the interaction neighborhood (i.e. within the area within distance $R$ from the cell surface) is then given by:

$$r = \frac{\left(R(R+w)(l-w) + \frac{4}{3}\left(\left(R+\frac{w}{2}\right)^3 - \left(\frac{w}{2}\right)^3\right)\right) \cdot \rho}{(l-w)(\frac{w}{2})^2 + \frac{w^3}{6}} \tag{22}$$

### 2 Predicting community level dynamics from local rules using pair-approximation—Model Derivation

We model a community consisting of two interacting cell types, $A$ and $B$, using evolutionary graph theory. In contrast to previous work, we allow the interaction neighborhoods to be different for the two types. As a result interactions are not symmetric, and we need to use directed graphs. We extend traditional pair approximation to find analytical expressions for the dynamics and the equilibrium state of the community. Pair approximation assumes that the spatial arrangements of any system can be described by only tracking pairwise correlations. As a consequence, the system can be fully described by tracking the number of all possible $(X \leftarrow Y)$ pairs. Pair approximation neglects any stochastic effects arising from finite populations and from finite numbers of cells in the interaction neighborhood.

#### 2.1 Definitions

We consider two cell types, $A$ and $B$, and use the notation $, Y \in \{A, B\}$ to indicate cell types. Cells are placed on the vertices of a directed graph. The total number of cells, $N$, is constant in time. At any given time, there are $N_A(t)$ and $N_B(t)$ cells of type $A$ and $B$, respectively. For notational simplicity, we will not explicitly show time dependence in our variables and simply write $N_A$ and $N_B$. Unless explicitly noted otherwise, all variable are time-dependent.

Interactions are encoded by directed links; in our notation $X \leftarrow Y$ indicates that focal cell $X$ interacts with neighboring cell $Y$, i.e. cell $X$ obtains metabolites from cell $Y$. All type $A$ cells interact with $r_A$ individuals, and we call the set of all neighbors with which a given $A$ cell interacts the small interaction neighborhood $I_A$. All type $B$ cells interact with $r_B$ individuals, and we call the set of all neighbors with which a given $B$ cell interacts the large interaction neighborhood $I_B$. Without loss of generality, we assume that type $A$ has the smallest interaction neighborhood, i.e. $r_A \leq r_B$. We also assume that the small interaction neighborhood is contained within the large one: $I_A \subseteq I_B$.

Because the neighborhood sizes of the two types are different, links are not symmetric, i.e. if there is a link $X \leftarrow Y$ between two cells, this does not imply that there is also a link $Y \leftarrow X$. The number of $Y \leftarrow X$ links is given by $N_{Y \leftarrow X}$. The total number of links in the system $L$ varies over time, because type $A$ and $B$ have different neighborhood sizes and the relative frequency of $A$ and $B$ changes in time:

$$L = N_{A \leftarrow A} + N_{A \leftarrow B} + N_{B \leftarrow A} + N_{B \leftarrow B} \tag{23}$$

Cells can place offspring within the replication neighborhood; the set of all neighbors that can be replaced is denoted by $I_R$, and counts $r_R$ cells. For mathematical tractability, we assume that the replication neighborhood is identical to smallest interaction neighborhood, i.e. $I_R = I_A$.

For each focal cell $X$, independently of its type, we can define two additional neighborhoods:

1) the small neighborhood $S(X, r_A)$ which consists of $r_A$ cells. This is the set of cells that can be replaced by the focal cell $X$ ($S(X, r_A) = I_R$). When we consider a focal cell of type $A$, this set of cells consists of all cells with which the $A$ focal cell interacts ($S(A, r_A) = I_A$). 2) the large neighborhood $S(X, r_B)$ which consists of $r_B$ cells. When we consider a focal cell of type $B$, this set of cells consists of all cells with which the $B$ focal cell interacts ($S(B, r_B) = I_B$). We assume $S(X, r_A) \subseteq S(X, r_B)$.

Throughout, we will use capital $N$ to denote numbers within the entire system, and small $n$ to denote numbers corresponding to the local neighborhood of a focal cell. For example, $N_A$ is the total number of $A$ cells in the system, while $n_A$ is the number of $A$ cells in the neighborhood of the focal cell.

#### 2.2 Pair-approximation

The system can be described by tracking the number of pairwise links $N_{X\leftarrow Y}$. We can express the total number of type $A$ and $B$ cells as function of $N_{X\leftarrow Y}$:

$$N_A = \frac{N_{A\leftarrow A} + N_{A\leftarrow B}}{r_A}, \quad N_B = \frac{N_{B\leftarrow A} + N_{B\leftarrow B}}{r_B} \tag{24}$$

each type $A$ focal cell has $r_A$ incoming links, which originate either from a type $A$ neighbor ($N_{A\leftarrow A}$ times) or from a type $B$ neighbor ($N_{A\leftarrow B}$ times). The number of $A$ cells $N_A$ can be found dividing the total number of incoming links over all $A$ cells by the number of incoming links per single $A$ cell. A similar reasoning allows to calculate the number of $B$ cells.

Because the total number of cells in the system ($N = N_A + N_B$) is constant, it follows from Eq. 24 that:

$$\frac{N_{A\leftarrow A} + N_{A\leftarrow B}}{r_A} + \frac{N_{B\leftarrow A} + N_{B\leftarrow B}}{r_B} = N = \text{constant} \tag{25}$$

We thus have three independent variables: one of the four quantities $N_{X\leftarrow Y}$ can be expressed as function of the other three.

To describe the composition of the system, we need the probability that a random cell we pick in the system is of either type $A$ or $B$. These probability, $P(A)$ and $P(B)$, follow directly from Eq. 24:

$$P(A) = \frac{N_{A\leftarrow A} + N_{A\leftarrow B}}{r_A N}, \quad P(B) = \frac{N_{B\leftarrow A} + N_{B\leftarrow B}}{r_B N} = 1 - P(A) \tag{26}$$

Moreover, we need to know the conditional probabilities that describe the local neighborhood of a cell. We define $P(Y|X, r_A)$ as the probability to find a neighbor of type $Y$ within the small neighborhood (i.e. in the set $S(X, r_A)$), given that the focal cell is of type $X$. Likewise, $P(Y|X, r_B)$ is the probability to find a neighbor of type $Y$ within the large neighborhood (i.e. in the set $S(X, r_B)$), given that the focal cell is of type $X$.

The conditional probabilities $P(Y|A, r_A)$ describe the average composition of the small neighborhood surrounding any type $A$ focal cell. By definition, the small neighborhood is identical to the interaction neighborhood of a type $A$ cell, $S(A, r_A) = I_A$. All cells in this small neighborhood, and only these cells, interact with the focal $A$ cell. All these cells are thus connected to the focal cells by $A \leftarrow Y$ links. From the number of $A \leftarrow Y$ links, we can directly calculate the conditional probabilities $P(Y|A, r_A)$:

$$P(A|A, r_A) = \frac{N_{A\leftarrow A}}{N_{A\leftarrow A} + N_{A\leftarrow B}}, \quad P(B|A, r_A) = 1 - P(A|A, r_A)$$

The equation for $P(A|A, r_A)$ can be intuitively understood as follows: the probability of finding a $A$ neighbor within the interaction neighborhood of an $A$ focal cell is simply the fraction of incoming links that start from $A$ cells.

Likewise, we can directly calculate the conditional probabilities $P(Y|B, r_B)$ from the number of $B \leftarrow Y$ links, because the large neighborhood is identical to the interaction neighborhood of type $B$, i.e. $S(B, r_B) = I_B$. We thus find:

$$P(A|B, r_B) = \frac{N_{B\leftarrow A}}{N_{B\leftarrow A} + N_{B\leftarrow B}}, \quad P(B|B, r_B) = 1 - P(A|B, r_B)$$

In contrast, the conditional probabilities $P(Y|A, r_B)$ and $P(Y|B, r_A)$ cannot be directly calculated from the pairwise links. $P(Y|A, r_B)$ describes the composition of the large neighborhood surrounding a type $A$ focal cell. This neighborhood can be larger than the interaction neighborhood of type $A$, and as a result the focal cell does not necessarily have incoming links from all cells in the large neighborhood. In other words, the set of cells in the interaction neighborhood of the type $A$ focal cell is only a subset of all cells in the large neighborhood: $I_A \subseteq S(A, r_B)$. $P(Y|B, r_a)$ describes the composition of the small neighborhood surrounding a type $B$ focal cell. The small neighborhood can be smaller than the interaction neighborhood of the type $B$ focal cell, i.e. $S(B, r_A) \subseteq I_B$. As a result, links to the type $B$ focal cell are not exclusive to the small neighborhood.

We can find the conditional probabilities $P(Y|A, r_B)$ and $P(Y|B, r_A)$ using a requirement of self-consistency. Suppose that we want to calculate the average number of type $A$ cells in the community. We can calculate this number directly from the global probability of finding a type $A$ cell as $N_A = P(A)N$. Alternatively, we can visit each cell in the community and calculate the average number of type $A$ neighbors it has. We will first do this considering the large $S(X, r_B)$ neighborhood for each cell. In $P(A)N$ cases the focal cell is of type $A$, which has $P(A|A, r_B)r_B$ type $A$ neighbors. In $P(B)N$ cases the focal cell is of type $B$, which has $P(A|B, r_B)r_B$ type $A$ neighbors. Each neighboring cell is counted $r_B$ times (it is part of the large $S(X, r_B)$ neighborhood of $r_B$ focal cells). Summing these two numbers and dividing by $r_B$ to correct for this multiple counting then gives the total number of $A$ cells in the system:

$$N_A = \frac{P(A)N \cdot P(A|A, r_B)r_B + P(B)N \cdot P(A|B, r_B)r_B}{r_B} = N\left(P(A) \cdot P(A|A, r_B) + P(B) \cdot P(A|B, r_B)\right)$$

Self-consistency requires that both methods give the same estimate for the average number of $A$ cells:

$$P(A)N = N\left(P(A) \cdot P(A|A, r_B) + P(B) \cdot P(A|B, r_B)\right)$$

From this we can thus find an expression for $P(A|A, r_B)$ as function of $P(A|B, r_B)$:

$$P(A|A, r_B) = 1 - \frac{P(B)}{P(A)} P(A|B, r_B)$$

We can repeat this procedure and count the number of $A$ and $B$ cells in the neighborhood of any $X$ cell, both using the small $S(X, r_A)$ and large $S(X, r_B)$ neighborhoods, to find expressions for the other undefined conditional probabilities $P(Y|A, r_B)$ and $P(Y|B, r_a)$. We can fully describe the local arrangements of the spatial system using the following set of conditional probabilities:

$$\begin{aligned}
P(A|A, r_A) &= \frac{N_{A\leftarrow A}}{N_{A\leftarrow A} + N_{A\leftarrow B}}, & P(A|A, r_B) &= 1 - \frac{P(B)}{P(A)} \cdot P(A|B, r_B) \\
P(B|A, r_A) &= \frac{N_{A\leftarrow B}}{N_{A\leftarrow A} + N_{A\leftarrow B}}, & P(B|A, r_B) &= \frac{P(B)}{P(A)} \cdot P(A|B, r_B) \\
P(A|B, r_A) &= \frac{P(A)}{P(B)} \cdot P(B|A, r_A), & P(A|B, r_B) &= \frac{N_{B\leftarrow A}}{N_{B\leftarrow A} + N_{B\leftarrow B}} \\
P(B|B, r_A) &= 1 - \frac{P(A)}{P(B)} \cdot P(B|A, r_A), & P(B|B, r_B) &= \frac{N_{B\leftarrow B}}{N_{B\leftarrow A} + N_{B\leftarrow B}}
\end{aligned} \tag{27}$$

#### 2.3 Cross-feeding communities

We will first use pair-approximation to derive predictions for the community-level properties of cross-feeding communities. In the next section we will generalize these results to any arbitrary community of two interacting cell types.

##### 2.3.1 Dynamical equations for community-level dynamics

We assume for now that growth rates depend linearly on the frequency of the partner type within the interaction neighborhood (we will later consider a more general growth function):

$$\mu_A(n_B) = \frac{n_B}{r_A} \cdot \hat{\mu}_A, \quad \mu_B(n_A) = \frac{n_A}{r_B} \cdot \hat{\mu}_B \tag{28}$$

where $n_B$ is the number of type $B$ cells within the interaction neighborhood of type $A$ cell, $n_A$ is the number of type $A$ cells within the interaction neighborhood of a type $B$ cell, $\hat{\mu}_A$ and $\hat{\mu}_B$ are the maximum growth rates of type $A$ and $B$, respectively, and $\langle\mu\rangle$ is the average growth rate of the community, which is given by:

$$\langle \mu \rangle = P(A) \cdot P(B|A, r_A) \cdot \hat{\mu}_A + (1 - P(A)) \cdot P(A|B, r_B) \cdot \hat{\mu}_B$$

Two events can change the number of links: a type $A$ cell reproduces and replaces a $B$ neighbor, with rate $T^+$, or a type $B$ cell reproduces and replaces an $A$ neighbor, with rate $T^-$. During a $T^+$ event the number of type $A$ cells thus increases by one, and during a $T^-$ event it decreases by one. To calculate rate $T^+$ we need to consider all events were an $A$ cell replaces a $B$ cell. A type $A$ cell can have $0 \leq n_B < r_A$ type $B$ neighbors. The probability of finding an $A$ cells with $n_B$ B neighbors is given by:

$$P(A) \cdot P(A|A, r_A)^{r_A - n_B} \cdot P(B|A, r_A)^{n_B} \cdot \binom{r_A}{n_B}$$

The probability that this cell reproduces is proportional to its growth rate, $\mu_A(n_B)$ relative to the average growth rate of the community as a whole $\langle \mu \rangle$:

$$\frac{n_B}{r_A} \cdot \frac{\hat{\mu}_A}{\langle \mu \rangle}$$

And the probability that the resulting offspring replaces a type $B$ cell is given by:

$$\frac{n_B}{r_A}$$

The rate $T^+$ can be found by multiplying these probabilities and summing over all possible number of $n_B$ neighbors:

$$T^+ = \sum_{n_B=0}^{r_A} P(A) \cdot P(A|A, r_A)^{r_A - n_B} \cdot P(B|A, r_A)^{n_B} \cdot \binom{r_A}{n_B} \cdot \frac{n_B}{r_A} \cdot \frac{\hat{\mu}_A}{\langle \mu \rangle} \cdot \frac{n_B}{r_A}$$

Performing the summation we find:

$$T^+ = P(A) \cdot P(B|A, r_A) \cdot \frac{1 + P(B|A, r_A)(r_A - 1)}{r_A} \frac{\hat{\mu}_A}{\langle \mu \rangle} \tag{29}$$

This equation can intuitively be understood as follows: the first factor gives the probability of choosing a type $A$ focal cell, the second the probability of choosing a type $B$ neighbor, and the third the probability that a type $A$ cell with at least one type $B$ neighbor reproduces. The term $1 + P(B|A, r_A)(r_A - 1)$ represents the expected number of $B$ neighbors, given that we know for sure that one neighbor is of type $B$ (the one that will be replaced by the offspring of $A$).

The rate $T^-$ can be found in a similar way, however we need to take into account that the replication neighborhood of type $B$ is smaller that its interaction neighborhood (i.e. $I_R \subseteq I_B$). The probability that the offspring of a $B$ cell replaces an $A$ neighbor is given by:

$$\frac{n_A}{r_B} \cdot \frac{P(A|B, r_A)}{P(A|B, r_B)}$$

the first factor is the probability of picking a type $A$ neighbor within the large neighborhood $I_B$, the second is the probability that an $A$ cell is part of the (small) replication neighborhood $I_R$, given that it is part of the (large) interaction neighborhood $I_B$. We thus find for $T^-$:

$$T^- = \sum_{n_A=0}^{r_B} (1 - P(A)) \cdot P(B|B, r_B)^{r_B - n_A} \cdot P(A|B, r_B)^{n_A} \cdot \binom{r_B}{n_A} \cdot \frac{n_A}{r_B} \cdot \frac{\hat{\mu}_B}{\langle\mu\rangle} \cdot \frac{n_A}{r_B} \cdot \frac{P(A|B, r_A)}{P(A|B, r_B)}$$

which sums to:

$$T^- = (1 - P(A)) \cdot P(A|B, r_B) \cdot \frac{1 + P(A|B, r_B)(r_B - 1)}{r_B} \cdot \frac{\hat{\mu}_B}{\langle\mu\rangle} \cdot \frac{P(A|B, r_A)}{P(A|B, r_B)} \tag{30}$$

When a $T^+$ or $T^-$ event happens the number of $X \leftarrow Y$ link changes by $\Delta^+_{XY}$ and $\Delta^-_{XY}$, respectively. We can use the conditional probabilities (Eq. 27) to calculate these quantities. Consider for example the case of a $T^+$ event, where an $A$ cell replaces a $B$ neighbor. The only links that change are the ones that start or end at the $B$ cell that will be replaced. To calculate the changes in links, we first have have to analyze the composition of the neighborhood of this $B$ cell. Using the conditional probabilities, we can calculate the expected number of $A$ and $B$ neighbors in both the small $S(B, r_A)$ and the large $S(B, r_B)$ neighborhood. We use our knowledge that the $B$ cell has at least one type $A$ neighbor (the cell that is about to reproduce) and find:

$$\begin{aligned}
n_A[S(B, r_A), T^+] &= 1 + (r_A - 1)P(A|B, r_A) \\
n_A[S(B, r_B), T^+] &= 1 + (r_B - 1)P(A|B, r_B) \\
n_B[S(B, r_A), T^+] &= (r_A - 1)P(B|B, r_A) \\
n_B[S(B, r_B), T^+] &= (r_B - 1)P(B|B, r_B)
\end{aligned}$$

here $n[S(B, r_A), T^+]$ indicates that this is the number of neighbors in the context of a small $S(B, r_A)$ neighborhood during a $T^+$ event.

These equations allow us to count the change of $B \leftarrow A$ links during a $T^+$ event. Before the replacement, all $B \leftarrow A$ links correspond to incoming links that start at an $A$ neighbor (within the large $S(B, r_B)$ neighborhood) that end at the central $B$ cell. We thus need to consider the number of $A$ neighbors within the large $S(B, r_B)$ neighborhood. In this neighborhood, we find $1 + (r_B - 1)P(A|B, r_B)$ $B \leftarrow A$ links before the replacement. After the replacement, $B \leftarrow A$ links corresponds to links that start at the newborn $A$ cell in the center and end at a $B$ neighbor (within the large $S(A, r_B)$ neighborhood). We thus need to consider the number of $B$ neighbors within the large $S(A, r_B)$ neighborhood. In this neighborhood, we find $(r_B - 1)P(B|B, r_B)$ $B \leftarrow A$ links[1]after the replacement. Taking the difference between the number of links before and after the replacement we thus find that:

$$\Delta^+_{BA} = (r_B - 1)P(B|B, r_B) - 1 - (r_B - 1)P(A|B, r_B)$$

Next, let's consider $B \leftarrow B$ links. These links are defined on the large $S(B, r_B)$ neighborhood. The central $B$ cell has $(r_B - 1)P(B|B, r_B)$ $B$ neighbors and for each neighbor there are two $B \leftarrow B$ links: one incoming (pointing to the central cell) and one outgoing. Before the replacement, there were thus $2(r_B - 1)P(B|B, r_B)$ $B \leftarrow B$ links. After the replacement, there are no $B \leftarrow B$ links, because the central cell is now of type $A$. We thus find:

$$\Delta^+_{BB} = -2(r_B - 1)P(B|B, r_B)$$

Using similar reasonings, we can derive the changes in the other links for both the $T^+$ and $T^-$ events:

$$\begin{aligned}
\Delta^+_{AA} &= 2 + 2(r_A - 1)P(A|B, r_A) \\
\Delta^+_{AB} &= (r_A - 1)P(B|B, r_A) - 1 - (r_A - 1)P(A|B, r_A) \\
\Delta^+_{BA} &= (r_B - 1)P(B|B, r_B) - 1 - (r_B - 1)P(A|B, r_B) \\
\Delta^+_{BB} &= -2(r_B - 1)P(B|B, r_B) \\
\Delta^-_{AA} &= -2(r_A - 1)P(A|A, r_A) \\
\Delta^-_{AB} &= (r_A - 1)P(A|A, r_A) - 1 - (r_A - 1)P(B|A, r_A) \\
\Delta^-_{BA} &= (r_B - 1)P(A|A, r_B) - 1 - (r_B - 1)P(B|A, r_B) \\
\Delta^-_{BB} &= 2 + 2(r_B - 1)P(B|A, r_B)
\end{aligned} \tag{31}$$

where we have used the following neighborhood properties in the context of the $T^-$ event:

$$\begin{aligned}
n_A[S(A, r_A), T^-] &= (r_A - 1)P(A|A, r_A) \\
n_A[S(A, r_B), T^-] &= (r_B - 1)P(A|A, r_B) \\
n_B[S(A, r_A), T^-] &= 1 + (r_A - 1)P(B|A, r_A) \\
n_B[S(A, r_B), T^-] &= 1 + (r_B - 1)P(B|A, r_B)
\end{aligned}$$

We can then write the dynamical equation for $N_{X \leftarrow Y}$ as:

$$\frac{dN_{X \leftarrow Y}}{dt} = T^+ \cdot \Delta^+_{XY} + T^- \cdot \Delta^-_{XY} \tag{32}$$

[1]Note that we use $P(B|B, r_B)$ even though after replication the cell at the center of the neighborhood is of type $A$. The reason is that the replication event only affects the central cell and not the wider neighborhood. Just before replication, a type $B$ cell was at the center of the neighborhood. The neighborhood is thus characterized by the conditional probabilities $P(Y|B, r)$ both directly before and directly after replication.

We now have all the information to solve the dynamical equations 32: by inserting the rates as given in Eq. 29 and 30, the changes in links given by 31, and the conditional probabilities given by 27. The system of four coupled differential equations can be solved numerically, or we can solve for the steady state analytically.

Before solving the dynamical equations, we change variables from $N_{X \leftarrow Y}$ to $P(A)$, $P(B|A, r_A)$, and $P(A|B, r_B)$:

$$P(A) = \frac{N_{A \leftarrow A} + N_{A \leftarrow B}}{rA \cdot N}, \quad P(B|A, r_A) = \frac{N_{A \leftarrow B}}{N_{A \leftarrow B} + N_{A \leftarrow A}}, \quad P(A|B, r_B) = \frac{N_{B \leftarrow A}}{N_{B \leftarrow A} + N_{B \leftarrow B}} \tag{33}$$

These new variables have direct biological meaning: the first corresponds to the global frequency of type $A$, the second corresponds to the local frequency of type $B$ in the interaction neighborhood of type $A$, and the last corresponds to the local frequency of type $A$ in the interaction neighborhood of type $B$.

Using Eq. 32 we can then write the dynamical equations for these variables:

$$\frac{dP(A)}{dt} = \frac{T^{+}\left(\Delta^{+}_{AA} + \Delta^{+}_{AB}\right) + T^{-}\left(\Delta^{-}_{AA} + \Delta^{-}_{AB}\right)}{N \cdot rA} \tag{34}$$

$$\frac{dP(B|A, r_A)}{dt} = -\frac{P(B|A, r_A)\Big(T^{-}\left(\Delta^{-}_{AA} + \Delta^{-}_{AB}\right) + T^{+}\left(\Delta^{+}_{AA} + \Delta^{+}_{AB}\right)\Big) - T^{-}\Delta^{-}_{AB} - T^{+}\Delta^{+}_{AB}}{N \cdot r_A \cdot P(A)} \tag{35}$$

$$\frac{dP(A|B, r_B)}{dt} = -\frac{P(A|B, r_B)\Big(T^{-}\left(\Delta^{-}_{BA} + \Delta^{-}_{BB}\right) + T^{+}\left(\Delta^{+}_{BA} + \Delta^{+}_{BB}\right)\Big) - T^{-}\Delta^{-}_{BA} - T^{+}\Delta^{+}_{BA}}{N \cdot r_B \cdot (1 - P(A))} \tag{36}$$

Where $T^{+}$ is given by Eq. 29, $T^{-}$ by Eq. 30, $\Delta^{+}_{XY}$ and $\Delta^{-}_{XY}$ by Eq. 31, and where the conditional probabilities (Eq. 27) can be expressed in terms of our new variables as:

$$\begin{aligned}
P(A|A, r_A) &= 1 - P(B|A, r_A), & P(A|A, r_B) &= 1 - \frac{1 - P(A)}{P(A)} \cdot P(A|B, r_B) \\
P(B|A, r_A) &= P(B|A, r_A), & P(B|A, r_B) &= \frac{1 - P(A)}{P(A)} \cdot P(A|B, r_B) \\
P(A|B, r_A) &= \frac{P(A)}{1 - P(A)} \cdot P(B|A, r_A), & P(A|B, r_B) &= P(A|B, r_B) \\
P(B|B, r_A) &= 1 - \frac{P(A)}{1 - P(A)} \cdot P(B|A, r_A), & P(B|B, r_B) &= 1 - P(A|B, r_B)
\end{aligned} \tag{37}$$

We can solve the temporal dynamics of these equation numerically.

##### 2.3.2 Steady state properties of cross-feeding communities

The equilibrium state can be found by setting Eq. 34-36 to zero and solving for $P(A)$, $P(B|A, r_A)$, and $P(A|B, r_B)$. We then find:

$$P(A) = \frac{\hat{\mu}_A \cdot \frac{r_A - 2}{r_A} + \left(\frac{\hat{\mu}_A}{r_A} - \frac{\hat{\mu}_B}{r_B}\right)}{\hat{\mu}_A \cdot \frac{r_A - 2}{r_A} + \hat{\mu}_B \cdot \frac{r_B - 2}{r_B}} \tag{38}$$

$$P(B|A, r_A) = \frac{r_A - 2}{r_A - 1} \cdot (1 - P(A)) \tag{39}$$

$$P(A|B, r_B) = \frac{r_B - 2}{r_B - 1} \cdot P(A) \tag{40}$$

$P(A)$ is the fraction of type A cells (i.e. it is a probability) and it thus has to satisfy $0 < P(A) < 1$. However, Eq. 38 can predict values for $P(A)$ which exceed these bounds. When Eq. 38 predicts values below 0 or above one this means that the two types cannot stably coexists.

In contrast, in a well mixed system, the growth rate of a cell only depends on the global frequency of the other cell type, from Eq. 28 it thus follows that:

$$\mu_A(P(A)) = (1 - P(A))\hat{\mu}_A, \qquad \mu_B(P(A)) = P(A)\hat{\mu}_B \tag{41}$$

At equilibrium the growth rate of both cell types is equal, which yields:

$$P(A)_{WM} = \frac{\hat{\mu}_A}{\hat{\mu}_A + \hat{\mu}_B} \tag{42}$$

#### 2.4 Arbitrary communities of two interacting cell types

We can generalize our findings to any system of two cell types that interact with a defined number of neighboring cells. When the growth function is linear we can find an analytical solution for the steady state properties of the community; for non-linear growth functions we can solve the dynamics numerically.

##### 2.4.1 Linear growth function

We consider the most general form of a linear growth function:

$$\begin{aligned} \mu_A(n_A, n_B) &= a_1 + a_2 \cdot n_A + a_3 \cdot n_B \\ \mu_B(n_A, n_B) &= b_1 + b_2 \cdot n_B + b_3 \cdot n_A \end{aligned} \tag{43}$$

here $a_1$ and $b_1$ represent the growth rate of type $A$ and $B$ cell when they are alone, $a_2$ and $b_2$ represent the increase in growth rate of type $A$ and $B$ cell when they are completely surrounded by their own type, and $a_3$ and $b_3$ represent the increase in growth rate of type $A$ and $B$ cell when they are completely surrounded by the other type. Our previous growth function (Eq. 28) can be recovered by setting $a_1, a_2, b_1, b_2 = 0$, $a_3 = \frac{\hat{\mu}_A}{r_A}$, and $b_3 = \frac{\hat{\mu}_B}{r_B}$. Because we assumed a birth-death process, it is important that the growth functions are positive for all possible neighborhood compositions.

For this growth function, the rates $T^+$ and $T^-$ are given by:

$$\begin{aligned} T^+ =& P(A) \cdot P(B|A, r_A) \cdot \qquad (44)\\ & \frac{1}{\langle\mu\rangle} \left(a_1 + a_2 \cdot P(A|A, r_A)(r_A - 1) + a_3 \cdot (1 + P(B|A, r_A)(r_A - 1))\right) \\ T^- =& (1 - P(A)) \cdot P(A|B, r_B) \cdot \qquad (45)\\ & \frac{1}{\langle\mu\rangle} \left(b_1 + b_2 \cdot P(B|B, r_B)(r_B - 1) + b_3 \cdot (1 + P(A|B, rB)(r_B - 1))\right) \cdot \frac{P(A|B, r_A)}{P(A|B, r_B)} \end{aligned}$$

Where the average community growth rate is given by:

$$\begin{aligned} \langle\mu\rangle =& P(A) \cdot (a_1 + a_2 \cdot P(A|A, r_A) r_A + a_3 \cdot P(B|A, r_A) r_A) + ... \\ & (1 - P(A)) \cdot (b_1 + b_2 \cdot P(B|B, r_B) r_B + b_3 \cdot P(A|B, r_B) r_B) \end{aligned} \qquad (46)$$

Solving Eq 34 for steady state, using the new rates for $T^+$ (Eq. 44) and $T^-$ (Eq. 44), we find:

$$P(A) = \frac{a_3(r_A - 2) - b_2(r_B - 2) + (a_1 + a_2 + a_3) - (b_1 + b_2 + b_3)}{(a_3 - a_2)(r_A - 2) + (b_3 - b_2)(r_B - 2)} \qquad (47)$$

As long as $0 < P(A) < 1$, the community has an equilibrium state where both types can coexist. We can also solve Eq. 35 and 36 at steady state and find that:

$$P(B|A, r_A) = \frac{r_A - 2}{r_A - 1} \cdot (1 - P(A)) \qquad (48)$$

$$P(A|B, r_B) = \frac{r_B - 2}{r_B - 1} \cdot P(A) \qquad (49)$$

The amount of clustering is thus the same, no matter what (linear) growth function is used.

##### 2.4.2 Non-linear growth functions

When the growth function is non-linear, in general it is not possible to find an analytical solution for the steady state. However, the ODE system given by Eq. 34-36 can always be solved numerically. For general, non-linear, growth functions $\mu_A(n_A, n_B)$ and $\mu_B(n_A, n_B)$ the following rates for $T^+$ and $T^-$ apply (it is in general not possible to solve these sums analytically):

$$T^+ = \sum_{n_B=0}^{r_A} P(A) \cdot P(A|A, r_A)^{r_A - n_B} \cdot P(B|A, r_A)^{n_B} \cdot \binom{r_A}{n_B} \cdot \frac{\mu_A(n_A, n_B)}{\langle\mu\rangle} \cdot \frac{n_B}{r_A} \tag{50}$$

$$T^- = \sum_{n_A=0}^{r_B} (1 - P(A)) \cdot P(B|B, r_B)^{r_B - n_A} \cdot P(A|B, r_B)^{n_A} \cdot \binom{r_B}{n_A} \cdot \frac{\mu_B(n_A, n_B)}{\langle\mu\rangle} \cdot \frac{n_A}{r_B} \cdot \frac{P(A|B, r_A)}{P(A|B, r_B)} \tag{51}$$

Our model can also be extended beyond a birth-death process, Eq. 34-35 would still hold, as long as appropriate rate functions are used for $T^+$ and $T^-$. The conditional probabilities as given by Eq. 37 and the expression for the changes in the number of links as given in 31 are independent of the replacement process and would thus also hold for e.g. a death-birth process.

### 3 Predicting community level dynamics from local rules using pair-approximation—Model Applications

#### 3.1 Cross-feeding interaction

Here we will consider how local interactions can decrease the average growth rate of cross-feeding communities. In a spatial system the growth rate of cells depends on the average local frequency of the partner type. The average community growth rate is thus given by:

$$\langle\mu\rangle = P(A) \cdot P(B|A, r_A) \cdot \hat{\mu}_A + (1 - P(A)) \cdot P(A|B, r_B) \cdot \hat{\mu}_B \tag{52}$$

This equation follows from the fact that a fraction of $P(A)$ cells is of type $A$, on average each of these cells as $P(B|A, r_A)r_A$ neighbors of type $B$, and thus grows at an average rate (see Eq. 28) of $P(B|A, r_A)\hat{\mu}_A$. Similarly, a fraction of $1 - P(A)$ cells is of type $B$, on average each of these cells as $P(A|B, r_B)r_B$ neighbors of type $A$, and thus grows at an average rate of $P(A|B, r_B)\hat{\mu}_B$.

In well mixed system the growth rate of cells depends on the global frequency of the partner type. The average community growth rate is thus given by:

$$\langle\mu\rangle = P(A) \cdot (1 - P(A)) \cdot \hat{\mu}_A + (1 - P(A)) \cdot P(A) \cdot \hat{\mu}_B \tag{53}$$

This equation follows from the fact that a fraction of $P(A)$ cells is of type $A$, on average each of these cells as $(1 - P(A))r_A$ neighbors of type $B$, and thus grows at an average rate (see Eq. 28) of $(1 - P(A))\hat{\mu}_A$. Similarly, a fraction of $1 - P(A)$ cells is of type B, on average each of these cells as $P(A)r_B$ neighbors of type $A$, and thus grows at an average rate of $P(A)\hat{\mu}_B$.

#### 3.2 Density dependent interaction

For some systems, we expect that growth is density and not frequency dependent. In this section we thus assume that growth rates increase linearly with the number, rather than the frequency, of neighbors of the other type:

$$\begin{aligned} \mu_A(n_A, n_B) &= \hat{\mu}_A \cdot n_B \\ \mu_B(n_A, n_B) &= \hat{\mu}_B \cdot n_A \end{aligned} \tag{54}$$

i.e. we use $a_1 = b_1 = a_2 = b_2 = 0$, $a_3 = \hat{\mu}_A$ and $b_3 = \hat{\mu}_B$. This linear growth function approximates well a more realistic Monod growth function, when the concentrations of the metabolite limiting growth are low in the environment, i.e. when they are below the saturation constant. Using Eq. 47, the steady state frequency is given by:

$$P(A) = \frac{\hat{\mu}_A(r_A - 2) + (\hat{\mu}_A - \hat{\mu}_B)}{\hat{\mu}_A(r_A - 2) + \hat{\mu}_B(r_B - 2)} \tag{55}$$

When $r_A, r_B \gg 1$ and $r_A \gg \frac{\hat{\mu}_B}{\hat{\mu}_A}$, this simplifies to:

$$P(A) = \frac{\hat{\mu}_A r_A}{\hat{\mu}_A r_A + \hat{\mu}_B r_B}$$

We thus see that the equilibrium frequency depends both on the strength of the interaction (given by $\hat{\mu}_A$ and $\hat{\mu}_B$) and the range of the interaction ($r_A$ and $r_B$); the type with the highest product $\hat{\mu} r$ will dominate the system.

#### 3.3 Growth inhibition

Our model can also be used to model communities with antagonistic interactions, where cells inhibits each others growth (e.g. by producing bacteriocins or antibiotics). We assume that the growth rate of each cell decreases linearly with the frequency of partner cell that produce a toxic substance, from a basal growth rate $\mu_0$ that is the same for both types:

$$\begin{aligned} \mu_A(n_A, n_B) &= \mu_0 - \delta_A \cdot \frac{n_B}{r_A} \\ \mu_B(n_A, n_B) &= \mu_0 - \delta_B \cdot \frac{n_A}{r_B} \end{aligned} \tag{56}$$

i.e. we use $a_1 = b_1 = \mu_0$, $a_2 = b_2 = 0$, $a_3 = -\frac{\delta_A}{r_A}$ and $b_3 = -\frac{\delta_B}{r_B}$. As we assume a birth-death process, it is essential that growth rates remain positive for all possible neighborhood compositions. We thus assume that $\delta_A, \delta_B < \mu_0$.

Using Eq. 47, the steady state frequency is given by:

$$P(A) = \frac{\delta_A \cdot \frac{r_A - 2}{r_A} + \left(\frac{\delta_A}{r_A} - \frac{\delta_B}{r_B}\right)}{\delta_A \cdot \frac{r_A - 2}{r_A} + \delta_B \cdot \frac{r_B - 2}{r_B}} \tag{57}$$

In the case of growth inhibition, the community as a whole grows better in space than in an equivalent well-mixed system. This can be seen from Eq. 48 and 48: in a spatial system the local frequency of the other cell type is lower than its global frequency. Because cells grow faster when there are fewer cells of the other type, cells in a spatial system grow faster. Unlike in the case for cross-feeding, where cells need the other type to grow well and thus grow worse when they interact with few neighbors, with growth inhibition cells actually grow better when they are surrounded by their own type and thus grow better when they interact with few neighbors.

##### 3.4 Application to an experimental cross-feeding community

We compared the prediction form our pair-approximation framework with data we previously obtained for an experimental synthetic cross-feeding community (1). In Table S1, we list all literature and measured parameter values used in this work. In Table S2, we compare the parameterization from experimental data with the parameterization from the key biophysical parameters. In Table S3, we compare the model predictions for the two parameterizations.

We previously found that the maximum growth rate predicted by Eq. 2 tends to be larger than the measured maximum growth rate (see Table S2 and ref. (1)). This is primarily because we assumed that all metabolites can be used for growth (i.e. we assume Monod kinetics), while in reality metabolites are also needed for cell maintenance. This maintenance reduces the amount of metabolites available for growth and thus reduces $\hat{\mu}$ (see (1) for more details). However, this equation can accurately predict the ratio of the maximum growth rates between the two cell types (see Table S2). From Eq. 38 it can be seen that our model only depends on the ratio of the maximum growth rates and not on their separate values, it is thus not needed to consider a more realistic growth function that includes maintenance costs.

#### 4 Supplementary Discussion

We modeled cell division using a "birth-death" process, were cells that divide replace one of their neighbors. In real systems, neighboring cells are not replaced, but pushed away. However, we believe that a birth-death process can model real systems because it retains two fundamental properties of these real systems. First, the reproduction rate of individuals is proportional to their growth rate; second, the offspring of an individual is placed close by in space.

The main difference is that in the model a random neighboring cell is removed to make place for the new cell, while in reality a random neighboring cell is pushed away. This generates a motion that is transmitted through the community, and as a result of this, another cell is pushed out to the edge. Cells at the edge are likely to be lost, for example because of shear stress of fluids flowing around the community (as is the case in our experimental growth chambers). The result of pushing a neighbor away is thus similar to the removal of a random cell from the system: in both cases a random cell is removed from the system, though this happens locally in the model and globally in reality.

Both in the model and in reality cell types are clustered in space i.e. they form patches. Removing a cell from the local neighborhood of a dividing cell is thus not exactly equivalent to removing a random cell from the system; the local neighbor is more likely to be of the same type as the dividing cell compared to a random cell. This could generate a discrepancy between the model and the real system.

We further assumed that the replication neighborhood is identical to the smaller interaction neighborhood ($r_R = r_A$). This assumption was needed for mathematical tractability, however we can easily relax it in our (individual based) cellular automaton simulations. We thus ran a batch of simulations were we set the replication neighborhood to be identical to the smallest interaction neighborhood ($r_R = r_A$) and a batch of simulations were we keep the replication neighborhood fixed at Moore neighborhood ($r_R = 8$). The results of these two batches closely match, even for relatively large interaction neighborhoods ($r_A = 120$, Fig S2), indicating that our simplifying assumption ($r_R = r_A$) does not affect our conclusions in any major way.

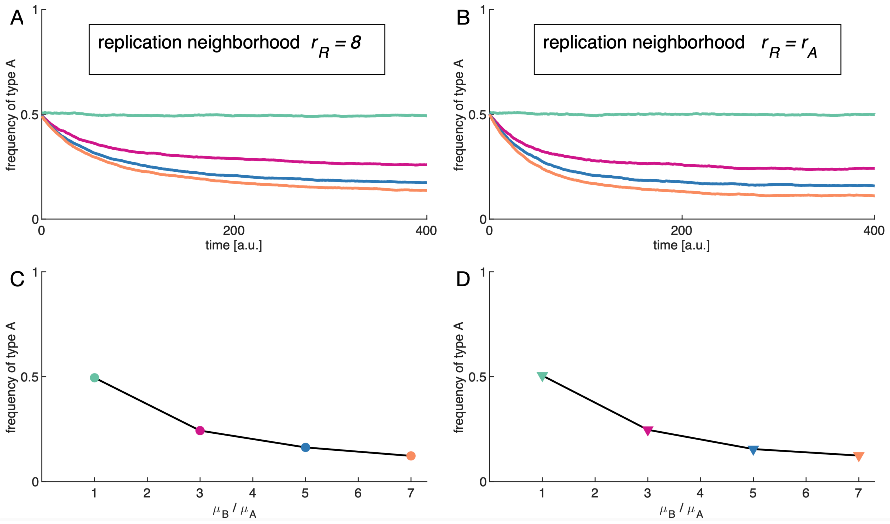

**Fig. S2:** Results of simulations are similar for a cellular automaton with replication neighborhood equal to the smallest interaction neighborhood ($r_R = r_A$) and for a cellular automaton with replication neighborhood equal to the Moore neighborhood ($r_R = 8$). (A) The fraction of type A is shown for four cellular automata simulations with $r_R = 8$ (Moore neighborhood). Each simulation is initialized with 50% of type A and 50% type B cells occupying a grid of 100x100. The four simulations have different values of $\mu_B/\mu_A$ and have $r_B = r_A = 120$. (B) Four cellular automata simulations with replication neighborhood $r_R = r_A$ and all other parameters as in panel A. Note that the steady state fraction of type A is reached slightly slower in panel A then in panel B. The cellular automata with $r_R = 8$ (C) reaches the same equilibrium as the cellular automata with $r_R = r_A$ (D) when having the same $\mu_B/\mu_A$. Each circle (or triangle) indicates the steady state of one of the simulations in panel A (or B).

### 5 Supplementary Tables

**Table S1: Parameters values used in study.** All parameters of the model are taken from literature or measured. Values are show as mean ± standard error of the mean. $l$, $w$, and $\rho$ were measured for 21 chambers, $l$, $w$ were averaged over all cells within a given chamber before averaging over all chambers. $\rho$ was estimated as the number of cells in the chamber divided by the total area occupied by cells. This area includes all intercellular free space, but excludes large empty areas in the chamber and was calculated by performing a morphological closing operation using a $1\mu m$ diameter on the segmented image. $^{a}$: the concentration is normalized with the Monod constant of the growth curve.

| Parameter | Description | Value | Source |
|---|---|---|---|
| $r^u_{\Delta P}$ | uptake rate of proline | $2.04\ 1/s$ | Literature (2) |
| $r^u_{\Delta T}$ | uptake rate of tryptophan | $24.05\ 1/s$ | Literature (3) |
| $D_{\Delta P}$ | diffusion rate of proline | $8.79 \cdot 10^2\ \mu m^2/s$ | Literature (4) |
| $D_{\Delta T}$ | diffusion rate of tryptophan | $6.59 \cdot 10^2\ \mu m^2/s$ | Literature (5) |
| $r^l_{\Delta P}$ | leakage rate proline | $1.59 \cdot 10^{-5}\ 1/s$ | Fitted (1) |
| $r^l_{\Delta T}$ | leakage rate tryptophan | $6.04 \cdot 10^{-7}\ 1/s$ | Fitted (1) |
| $R_{\Delta P}$ | interaction range of ΔP | $12.1 \pm 0.5\ \mu m$ | Measured (1) |
| $R_{\Delta T}$ | interaction range of ΔT | $3.2 \pm 0.4\ \mu m$ | Measured (1) |
| $\beta$ | constant of proportionality between growth and interaction range | 0.88 | Measured (1) |
| $\mathcal{I}^{\mathcal{C}}$ | normalized concentration of metabolite in producing cell$^a$ | 20 | Estimated (1) |
| $\mu^{wt}$ | growth on M9 media + 0.2% glucose | $1.29\ 1/h$ | Measured (1) |
| $\rho$ | volume density of cells | 0.65 | Measured (1) |
| $\rho_{2D}$ | area density of cells | $0.22 \pm 0.01$ cells/$\mu m^2$ | Measured |
| $l$ | average cell length | $5.2 \pm 0.1\ \mu m$ | Measured |
| $w$ | average cell width | $0.68 \pm 0.01\ \mu m$ | Measured |

**Table S2: Model parameterization.** We compare the model parameters for the parameterization based on experimental data and the one based on the biophysical model (Eq. 1 and 2). The biophysical model overestimates the maximum growth rates, however it can accurately predict the ratio of the maximum growth rates, and this is the only factor that matters in our model. For the cellular automaton simulations the interaction neighborhood was set to an extended Moore neighborhood with radius $d$ such that the number of cells in the Moore neighborhood $(2 \cdot d + 1)^2 - 1$ most closely matched the estimated number of neighbors $r$.

| **Parameter** | **Description** | **Value estimated from data** | **Value estimated from biophysical parameters** |
|---|---|---|---|
| $r_{\Delta P}$ | number of neighbors of ΔP | 129 cells | 130 cells |
| $r_{\Delta T}$ | number of neighbors of ΔT | 15 cells | 10 cells |
| $d_{\Delta P}$ | radius of extended Moore neighborhood of ΔP | 5 grid units (120 cells) | 5 grid units (120 cells) |
| $d_{\Delta T}$ | radius of extended Moore neighborhood of ΔT | 1 grid unit (8 cells) | 1 grid unit (8 cells) |
| $\hat{\mu}_{\Delta P}$ | maximum growth rate of ΔP | 0.52 $1/h$ | 0.77 $1/h$ |
| $\hat{\mu}_{\Delta T}$ | maximum growth rate of ΔT | 0.15 $1/h$ | 0.22 $1/h$ |
| $\frac{\hat{\mu}_{\Delta P}}{\hat{\mu}_{\Delta T}}$ | growth ratio | 3.5 | 3.6 |

**Table S3: Comparison of model parameterizations.** We compare the predictions from the model parameterized from experimental data with predictions from the model parameterized from the biophysical model (Eq. 1 and 2).

| **Variable** | **Predicted using parameterization from data** | **Predicted using parameterization from biophysical parameters** | **Measured value** |
|---|---|---|---|
| $P(\Delta T)$ | 0.21 | 0.20 | 0.19 |
| $\frac{P(\Delta T \mid \Delta P, r_{\Delta P})}{P(\Delta T)}$ | 0.99 | 0.99 | 0.87 |
| $\frac{P(\Delta P \mid \Delta T, r_{\Delta T})}{P(\Delta P)}$ | 0.93 | 0.89 | 0.85 |
| Community growth rate relative to well-mixed | 0.94 | 0.92 | 0.87 |
